# Multiomics reveals persistence of obesity-associated immune cell phenotypes in adipose tissue during weight loss and subsequent weight regain

**DOI:** 10.1101/2021.08.20.455954

**Authors:** Matthew A. Cottam, Heather L. Caslin, Nathan C. Winn, Alyssa H. Hasty

**Author notes:** These authors contributed equally to this work (Matthew A. Cottam, Heather L. Caslin). Corresponding author: Alyssa Hasty.

## Abstract

Most individuals do not maintain weight loss, and weight regain increases cardio-metabolic risk beyond that of obesity. Adipose inflammation directly contributes to insulin resistance; however, immune-related changes that occur with weight loss and weight regain are not well understood. Single cell RNA-sequencing was completed with CITE-sequencing and biological replicates to profile changes in murine immune subpopulations following obesity, weight loss, and weight cycling. Weight loss normalized glucose tolerance, however, type 2 immune cells did not repopulate adipose following weight loss. Many inflammatory populations persisted with weight loss and increased further following weight regain. Obesity drove T cell exhaustion and broad increases in antigen presentation, lipid handing, and inflammation that persisted with weight loss and weight cycling. This work provides critical groundwork for understanding the immunological causes of weight cycling-accelerated metabolic disease. Thus, we have created an open-access interactive portal for our processed data to improve accessibility for the research community.

## Introduction

Obesity affects more than 650 million adults worldwide and is associated with nearly every leading cause of death, including cardiovascular disease, diabetes, and several types of cancer^1, 2^. In lean adipose tissue (AT), regulatory type 2 immune cells contribute to tissue homeostasis^3, 4, 5, 6^. In contrast, weight gain promotes the infiltration of circulating immune cells, and tissue resident cells polarize towards a type 1 pro-inflammatory phenotype. These inflammatory changes contribute to metabolic dysfunction – promoting lipolysis, fibrosis and insulin resistance (IR).

Weight loss (WL) is known to improve metabolic outcomes associated with obesity. However, low success rates and failure to maintain lost weight are common, with recent studies reporting that most individuals (>60%) regain weight within a few years^7, 8, 9, 10^. Importantly, weight cycling (WC) – the repeated process of gaining and losing weight – further increases risk for developing cardiometabolic diseases^11, 12, 13^. We generated a model of WC using alternating high fat diet (HFD) and low fat diet (LFD) feeding^14^. These mice display worsened glucose tolerance compared with obese mice, despite similar body weight, fat mass, and total time on the HFD. As assessed by flow cytometry, metabolic dysfunction was associated with increases in AT T cell populations, but not macrophages, consistent with other models of WC^15^.

With obesity, cells can upregulate type 1 and type 2 transcriptional profiles and metabolic pathways simultaneously^16, 17^. Moreover, recent single-cell RNA-sequencing (scSeq) studies have revealed multiple distinct AT immune cell populations and subsets, such as lipid associated macrophages (LAMs), which are unique from traditional macrophage polarization states^17, 18, 19^. scSeq allows for the interpretation of changes to immune populations and unbiased gene expression across the entire immune landscape simultaneously. Further developments, such as Cellular Indexing of Transcriptomes and Epitopes by sequencing (CITE-seq) and cell hashing, offer opportunities to interrogate surface protein repertoire and include biological replicates during scSeq experiments^20, 21^.

To date, no studies have thoroughly characterized AT immune populations following WC by a comprehensive technique like scSeq, and many scSeq datasets are difficult to interrogate further without bioinformatics expertise. Therefore, the aims of this study were two-fold: 1) To comprehensively map the changes in adipose immune populations with obesity, WL, and WC and 2) To provide a freely accessible resource for hypothesis generation and testing for the scientific community in a novel dataset that spans four distinct physiological states of AT. Broadly, we found that obesity-associated inflammatory changes such as T cell exhaustion and macrophage lipid handling persist following WL and may contribute to WC-accelerated metabolic disease.

## Results

### Diet induced WC exacerbates glucose intolerance

We used 9-week bouts of HFD and LFD feeding to generate models of obesity, WL, and WC (**Fig 1A**)^14^. This model is robust, as shown across 6 cohorts (totaling 252 mice). Changes in weekly body mass and energy intake were tightly linked to prescribed diets (**Fig 1B &C**). Cumulative energy intake was not different between obese, WL, and WC groups (**Fig 1D**). After 26-weeks on diet, obese and WC mice had identical lean and fat mass, which was elevated compared to lean and WL mice that also had identical lean and fat mass (**Fig 1E**). Obese mice had impaired glucose tolerance by intraperitoneal glucose tolerance tests (ipGTT) compared to lean mice, which was further worsened by WC (**Fig 1F**). A small improvement in glucose tolerance was observed in WL mice compared to lean control mice that can be largely attributed to reduced peak plasma glucose levels. Subcutaneous AT (sAT) and liver weights were similar between lean vs. WL and obese vs. WC mice. However, WL mice had reduced epididymal (eAT) mass compared to lean mice, and WC mice had increased eAT mass compared to obese mice (**Fig 1G**). Immuno-fluorescence staining for perilipin-1, a lipid droplet membrane protein, revealed a slight reduction in lipid droplet diameter in eAT of WL compared to lean mice, while no difference was observed between obese and WC mice. (**Fig 1H & I**). The corresponding body mass, food intake, tissue mass, and ipGTT for the subset of 4 mice per group included in subsequent scSeq experiments is shown in **Fig S1**. Together, these data demonstrate that our mouse model provides a robust representation of WC-accelerated metabolic disease.

**Figure 1.**
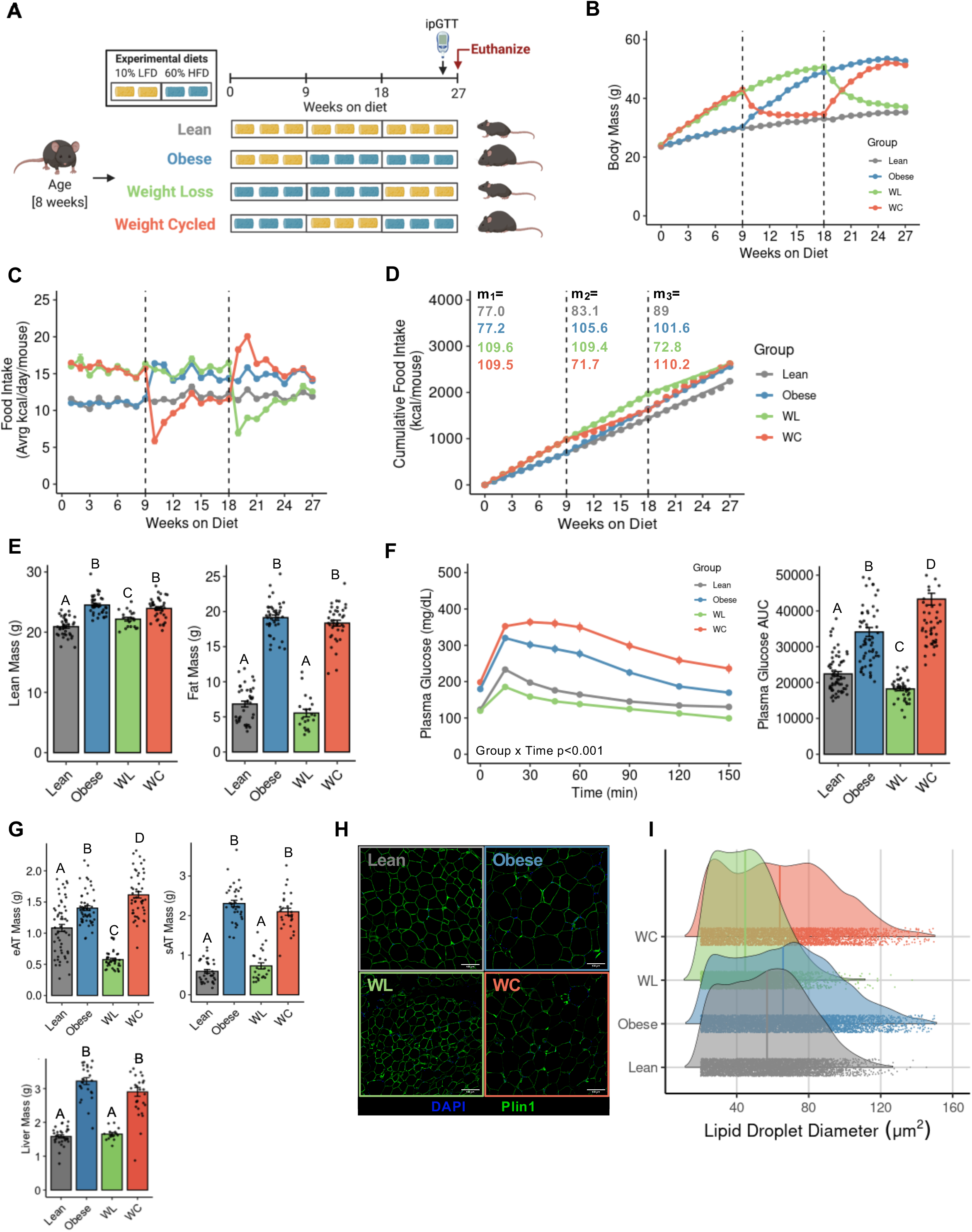
Mouse models of lean, obese, weight loss and weight cycling. (A) Schematic of dietary approaches to generate WL and WC mice. (B) Body mass over time measured weekly with diet switch indicated by dashed lines. (C) Food intake over time measured weekly. (D) Cumulative food intake measured throughout the duration of the studies with slope for each 9-week segment indicated. (E) Lean and fat mass measured by NMR. (F) Blood glucose during an ipGTT dosed at 1.5 g dextrose/kg lean mass one week prior to the end of the study and area under the curve for ipGTT. (G) Tissue mass at sacrifice. (H) Representative imaging of Perilipin-1 immunofluorescence for (I) lipid droplet size quantification with median and distribution for 5,000 cells per group. Pairwise two-tailed Student’s t-tests with Bonferroni correction for multiple comparisons were used to compare groups for body composition, tissue mass, and ipGTT AUC and two-way ANOVA was used to compare groups for ipGTT (groups indicated by different letters are significantly different, \**adj p* < 0.05. Data is plotted as mean ± SEM (n=39-62).

### Multimodal single-cell sequencing highlights the diversity of adipose immune cells

To profile the immune landscape across lean, obese, WL, and WC groups, we performed droplet based scSeq with CITE-seq, whereby oligo-conjugated antibodies specific for surface protein targets are simultaneously sequenced with endogenous mRNA. We also used cell hashing via antibodies conjugated with unique oligos targeting ubiquitous surface proteins to pool and index 4 biological replicates per group. Cells were isolated via collagenase digestion, CD45^+^ magnetic enrichment, and FACS for viability before preparation for sequencing using the 10X Chromium platform (**Fig 2A**). CITE-seq antibodies were validated by comparing sequenced protein expression levels to gene expression levels for matched targets (i.e. CD4 vs. *Cd4*), largely exclusive targets (i.e. CD4/*Cd4* vs. CD19/*Cd19*), and for commonly associated targets (i.e. CD4/*Cd4* vs. CD3ε/*Cd3e*) (**Fig S2**). Importantly, biological replicates are frequently ignored in scSeq experiments, and we sought to address this concern using cell hashing antibodies. Cell types were well represented in all biological replicates without any major outliers driving our interpretation and cell classification (**Fig S3**).

**Figure 2.**
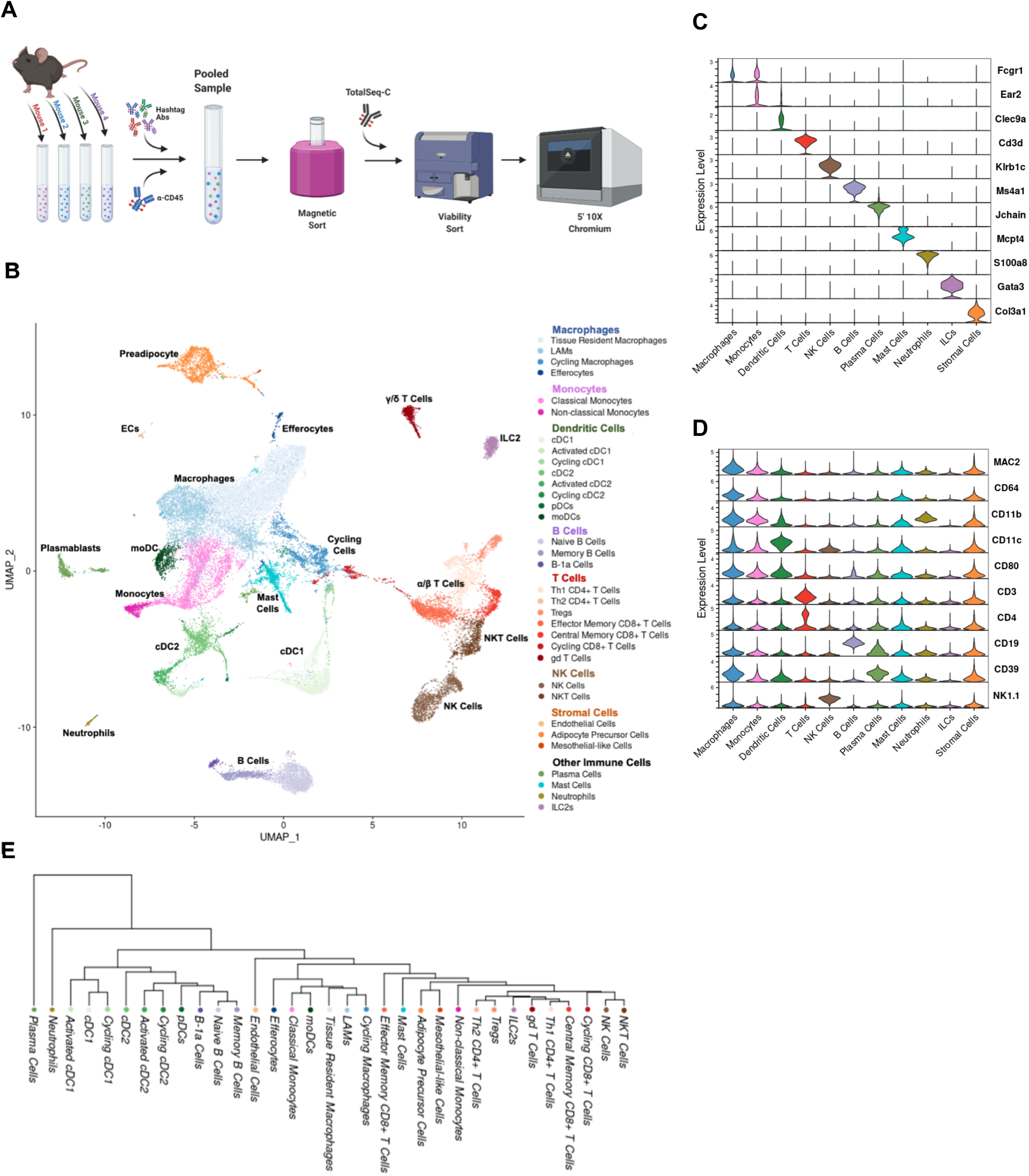
Adipose tissue immune cell populations observed by CITE-seq. (A) Schematic of CITE-seq approach. (B) Unbiased clustering of 33,322 single cells labeled broadly by cell type category and colored by high-resolution cell type identities. Selected markers of specific cell subsets based on (C) gene expression and (D) surface protein. (E) Phylogenetic tree of high-resolution cell type identities.

Across the four sample groups (16 mice), a total of 33,322 cells that met strict quality control metrics (see *Methods*) were retained and integrated. Large cell clusters generated using low resolution nearest-neighbor clustering were first annotated using differentially expressed protein and gene signatures. Further subcluster annotation was conducted by subsetting large cell type clusters. A heatmap of the top 5 genes associated with each celltype subcluster is shown in **Fig S4**. The resulting dimensional reduction with cells colored by subcluster is shown in **Fig 2B**. A subset of genes associated with specific subclusters are highlighted in **Fig 2C**. Antibody-based protein measurements for highly expressed surface proteins were also robust for cell type identification (**Fig 2D**). Finally, a dendrogram was produced using the 2,000 most highly variable genes to elucidate broad relationships between cell subclusters (**Fig 2E**). Cell cluster designations were confirmed by published gene markers and are used throughout the rest of this manuscript to interrogate group differences by diet.

### Obesity-associated immune cell phenotypes are confirmed by single cell sequencing

To validate our scSeq dataset, we compared cells isolated from obese mice to those from lean mice, as these differences are well documented. To explore how relative frequency of immune cells changed during obesity, we grouped cells into metacells based on UMAP proximity and compared abundance by diet group (**Fig 3A**). By linking metacells to original cell type designations, we observe subpopulations of T cells, dendritic cells (DCs), and macrophages that are differentially abundant in lean and obese eAT (**Fig 3B**), consistent with published literature^6, 22, 23^. Furthermore, metacells were reclassified by high-resolution subclusters to identify potential subpopulations of interest, such as regulatory T cells (T_regs_), type 2 innate lymphoid cells (ILC2s), and tissue resident macrophages (TRMs) – abundant in the lean state, and LAMs, effector memory CD8^+^ T cells (CD8^+^ T_EM_) and activated DCs – abundant in the obese state (**Fig 3C**). By tracing cells back to each individual mouse, we were also able to perform pairwise comparisons of cell number for cell subtypes classically associated with lean AT (**Fig 3D**) and obese AT (**Fig 3E**). These data validate our cell isolation strategy and cell type identification by confirming, with high fidelity, many of the previously established changes associated with lean and obese AT.

**Figure 3.**
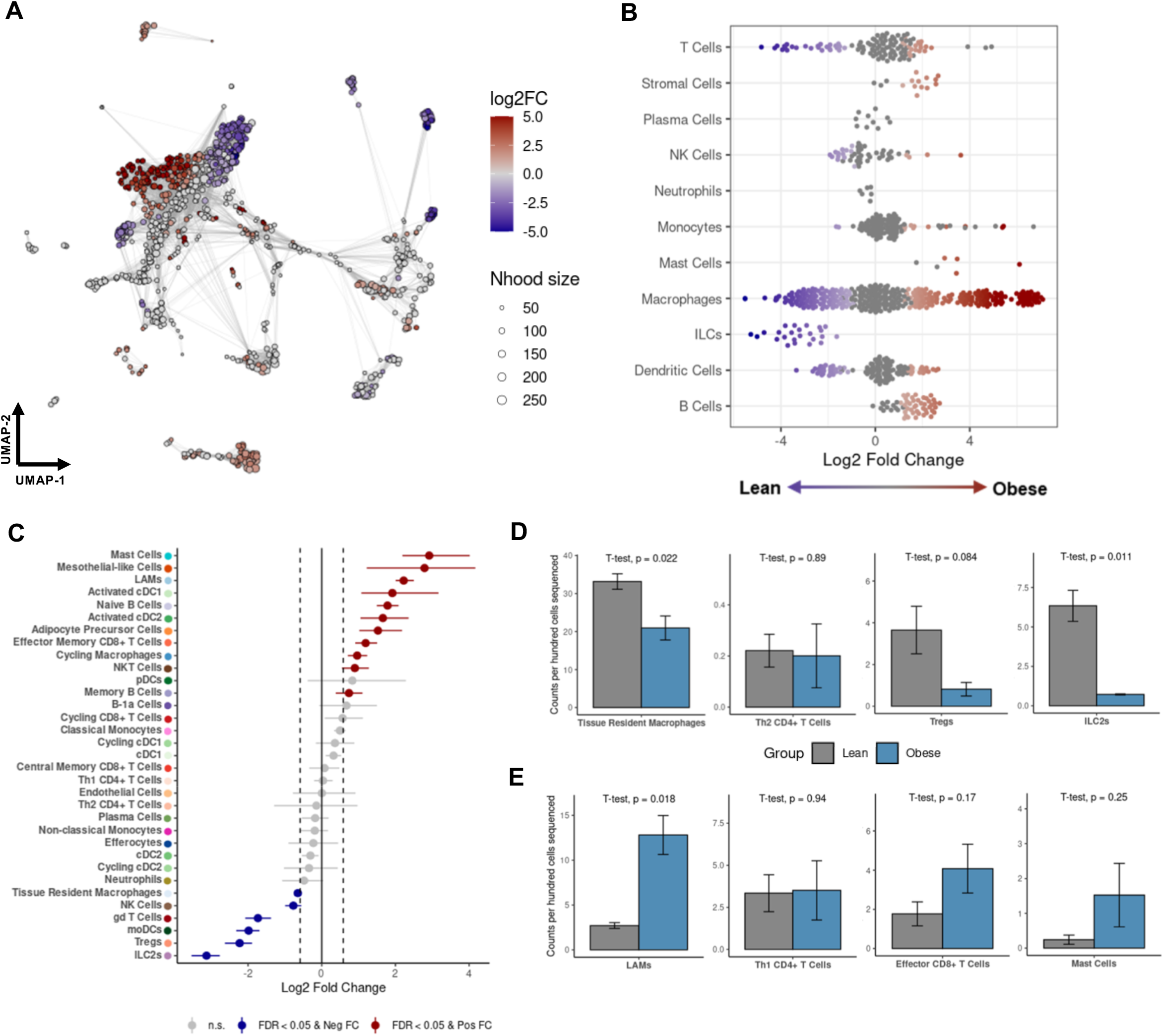
CITE-seq recapitulates obesity-associated immune cell changes in adipose tissue. (A) Differential abundance of metacells comparing cells from obese mice to lean mice. (B) Differential abundance of cell types in lean and obese mice. (C) Permutation testing of high-resolution clusters to calculate the proportional difference in lean and obese mice. (D & E) Counts per hundred cells sequenced for lean-associated immune cell subsets and obesity-associated immune cell subsets in lean and obese mice (mean ± SEM; n=4; two-tailed t-test with indicated *p* values).

### Obesity-associated T cell exhaustion persists after WL

Adaptive T-lymphocytes are important regulators and drivers of inflammation in AT. We explored AT T cells by scSeq (**Fig 4A**) and identified specific subsets of α/β-T Cells by expression of conventional T cell markers (*Cd3ε, Cd4, Cd8b1)*, phenotype markers (*Foxp3,Cxcr3, Ccr7, Sell*), and markers of cell cycling (*Stmn1, Pclaf*) (**Fig 4B**). A population of γ/δ-T cells was identified by expression of the common T cell delta chain (*Trdc*) and both natural killer (NK) and natural killer T cells (NKT) were identified by expression of killer lectin receptors *Klrb1b* and *Klrb1b* with and without expression of *Cd3ε*, respectively (**Fig S5**). Th1 CD4^+^ and all three CD8^+^ T cells subsets were not elevated in WC but CD8^+^ effector memory T cells were increased in AT from WL mice (**Fig 4C**). T_regs_ expressing *Foxp3* trended towards being reduced in proportion by obesity and remain low following WL and WC. Using differential expression, we observed that obesity not only affected T_reg_ number but also reduced the expression of *Il1rl1,* which codes for the IL-33 receptor, ST2 (**Fig 4D**). Changes in other IL-33 responsive cell types, such as ILC2s and mast cells were also observed following obesity, failed to recover with WL, and were exacerbated by WC (**Fig S6**).

**Figure 4.**
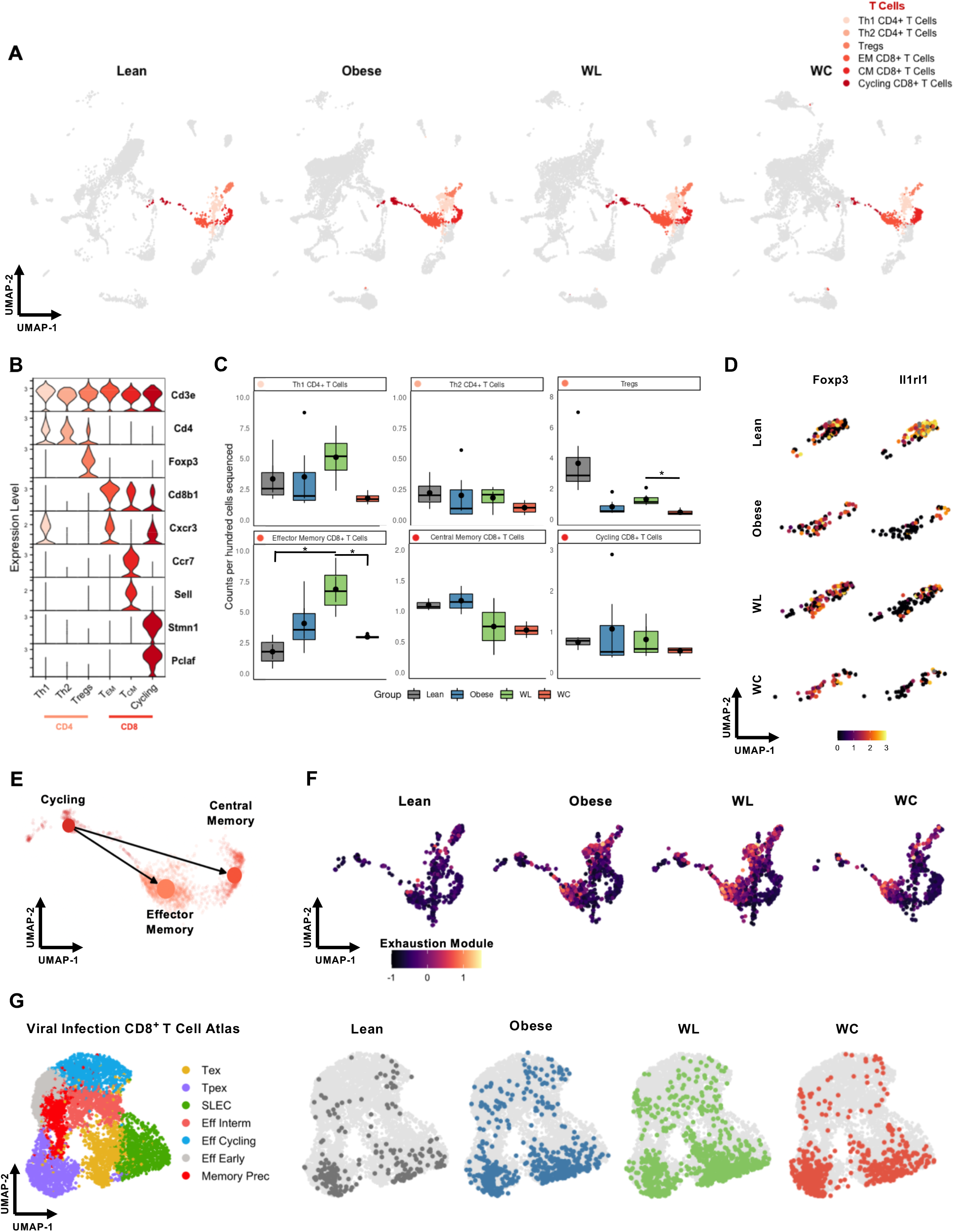
Adipose tissue T cells are retained and express markers of exhaustion following onset of obesity. (A) UMAP of T cell subclusters by diet group. (B) Expression of markers enriched in T cell subclusters. (C) Counts per hundred cells sequenced for α/β T cell subclusters (mean ± sem; n=4). (D) Expression of the T_reg_ marker *Foxp3* and *Il1rl1* within the T_reg_ subcluster across diet groups. (E) Predicted trajectory of CD8^+^ T cells by partition-based graph abstraction (PAGA) with directionality inferred by RNA velocity. (F) CD8^+^ T cells colored by an exhaustion module containing the features *Pdcd1, Tox, Tigit, Lag3, and Entpd1*. (G) CD8^+^ T cells plotted onto a viral infection CD8^+^ T cell reference atlas using ProjecTILs. Pairwise two-tailed t-tests with Bonferroni correction for multiple comparisons were used to compare groups for cell counts (*adj *p* < 0.05).

Because CD8^+^ T cells have the capacity to clonally expand in response to antigen, we used RNA velocity directed partition-based graph abstraction (PAGA) to explore whether cycling cells (marked by expression of *Pclaf, Stmn1,* and *Mki67*) are precursors for AT memory T cell populations. PAGA indicates that cycling CD8*^+^* T cells are upstream of other effector and memory populations in AT (**Fig 4E**). Differential expression across diet groups also identified that cycling and effector memory (T_EM_) CD8^+^ T cells express the activation/exhaustion signature gene *Pdcd1* (coding for PD-1). To further explore whether AT T cells contain an exhausted signature following obesity, we generated a module containing multiple established features of T cell exhaustion: *Pdcd1, Tox, Entpd1, Tigit,* and *Lag3*. We observed that obesity, WL, and WC were all enriched for our exhaustion module and that CD8^+^ T_EM_ were most associated with an exhausted phenotype (**Fig 4F**). T cell exhaustion is frequently studied in models of viral infection and in the tumor microenvironment, so we utilized ProjecTILs, a published scRNA-seq reference atlas^24^, to further confirm our exhaustion profile. We observed that cells captured in our experiments projected more accurately onto an LCMV chronic infection atlas (**Fig 4G**) than to a tumor-infiltrating lymphocyte (TIL) atlas (**Fig S7**). Concordant with our exhaustion module, more CD8^+^ T cells aligned with the LCMV exhausted precursor cells in obese, WL, and WC groups than in the lean control group.

### Monocytes are abundant in AT, but do not have an altered transcriptional profile in response to obesity, WL, or WC

The AT immune cell compartment contains a large population of both monocytes and DCs (**Fig 5A**). Monocytes were subclustered into classical (*Ly6c2*^+^*Ccr2^+^Cd14^+^*) and non-classical (*Ly6c2^−^Ear2^+^Cx3cr1^+^*) subsets (**Fig 5B**). Additionally, we observed that many classical monocytes expressed *Fcgr1* and that *Ace* was highly specific for non-classical monocytes in AT. We did not observe strong correlation between diet group and number of either classical or non-classical monocytes (**Fig 5C**). To investigate whether monocyte phenotype was different between diets, we aggregated counts and analyzed monocytes as a pseudobulk dataset whereby each unique mouse was treated as a separate sample. Using a likelihood ratio test through DESeq2, any genes showing changes in expression across diet groups were plotted into distinct patterns and pathway analysis was performed on each mutually exclusive gene set (**Fig 5D**). Pathway analysis suggests that genes which recover with WL are associated with protein and cholesterol synthesis and fatty acid metabolism, while those that fail to recover are associated with response to cellular stimuli and leukocyte differentiation.

**Figure 5.**
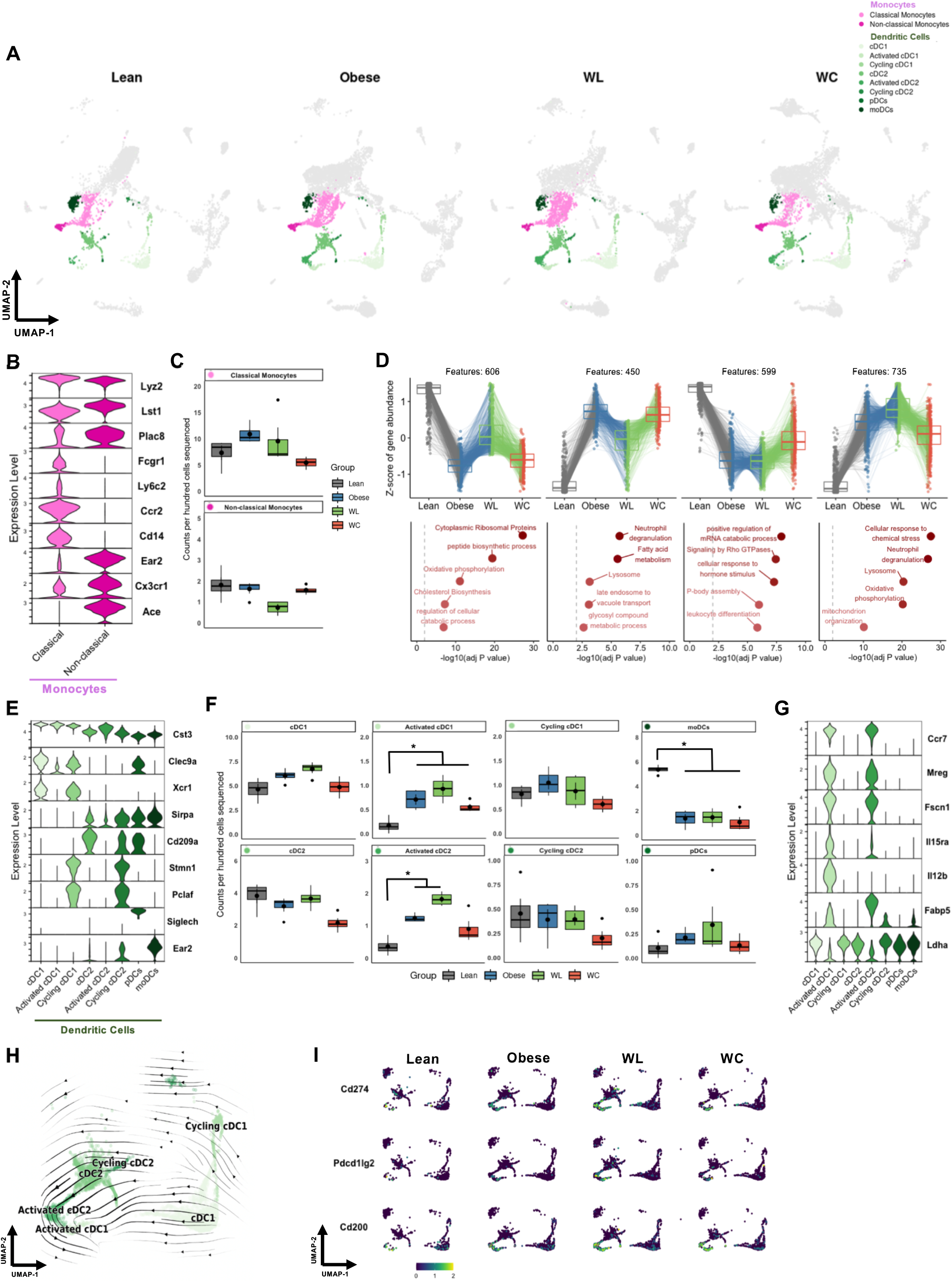
Monocytes and DCs are abundant in AT and DCs shift towards a mature, activated status. (A) Highlighted monocyte and DC subclusters. (B) Expression of genes associated with monocyte subclusters. (C) Monocyte counts per 100 cells sequenced by diet group (n=4; n.d.). (D) Patterns of mutually exclusive gene changes in monocytes across diet groups and the top 5 hits for pathway analysis for each pattern. (E) Expression of genes enriched in DC subsets. (F) Dendritic cell counts per 100 cells sequenced by diet group (mean ± sem; n=4). (G) Expression of genes associated with DC activation. (H) Embedding of RNA velocity displayed on the UMAP for cDC subsets. (I) Expression of immunoregulatory ligands Cd274, Pdcd1lg2, and CD200 in DCs by diet group. Pairwise two-tailed T-tests with Bonferroni correction for multiple comparisons were used to compare groups against the Lean reference group for cell counts (\**adj p* < 0.05).

### DCs shift towards an activated transcriptional signature with obesity and these signatures are retained with WL and WC

DCs, a class of professional antigen presenting cells (APCs), make up approximately 10-15% of captured AT immune cells. We classified DCs into two conventional DC (cDC) subsets, cDC1s (*Clec9a^+^Xcr1^+^*) and cDC2s (*Sirpa^+^Cd209^+^*), and further refined subclustering using markers of DC activation (*Ccr7*) and cell cycle induction (*Pclaf and Stmn1*) (**Fig 5E**). AT DCs were largely unchanged in relative proportion by diet condition with the exception of monocyte-derived DCs (moDCs), which were enriched in lean AT (**Fig 5F**). Activated subsets of DCs were elevated in obese AT and remained elevated upon WL and WC. Expression of *Ccr7* correlated positively with expression of *Fscn1* and *Mreg* and negatively with expression of cDC1 and cDC2 markers, indicating that activated DC subsets are mature (**Fig 5G**). *Il15ra*, which codes for the receptor that trans-presents IL-15 to NK and T cells to support homeostatic proliferation^25, 26^, was also correlated with *Ccr7*. RNA velocity was used to determine whether activated DCs were precursors or downstream of other cDC subsets and suggested that activated cDCs mature from other local cDC subsets (**Fig 5H**). Expression of *Ccr7* also negatively correlated with lactate dehydrogenase expression (*Ldha*), suggesting the potential for a shift in metabolite usage upon activation. Activated cDC1s, but not cDC2s, also expressed high levels of *Il12b* and may be primed to induce a local Th1 response within AT. The activated DC subset also expressed very high levels of key immunoregulatory ligands *Cd274, Pdcd1lg2,* and *Cd200* associated with coinhibitory T cell receptors **(Fig 5I).** Taken together, these data suggest that DCs in AT shift towards an activated immunoregulatory status with onset of obesity and that these features are retained long-term through WL and WC.

### Macrophage populations are highly adaptable to change in dietary status

Macrophages make up the largest proportion of immune cells in AT and are highly responsive to changes initiated by caloric excess. Dimensional reduction and clustering of macrophage subsets highlights the dramatic changes that occur during AT adaptation to HFD (**Fig 6A**). Common markers of tissue macrophages, *Lyz2*, *Cst3*, *Adgre1* (coding for F4/80), *Cd68,* and *Lgals3* (coding for MAC2) were highly expressed by all macrophage subclusters and robust protein expression for conventional AT macrophage markers CD64 and CD11b was observed (**Fig 6B**). TRMs were enriched for expression of *Klf4, Cbr2, and Stab1*, while LAMs highly express various genes associated with lipid interactions (*Trem2, Cd9, Lpl*) (**Fig 6C**). In addition, we identified a subset of cycling macrophages and another small subset of macrophages that expressed very high levels of *Saa3 and Slpi,* which are defining features of efferocytes^27^.

**Figure 6.**
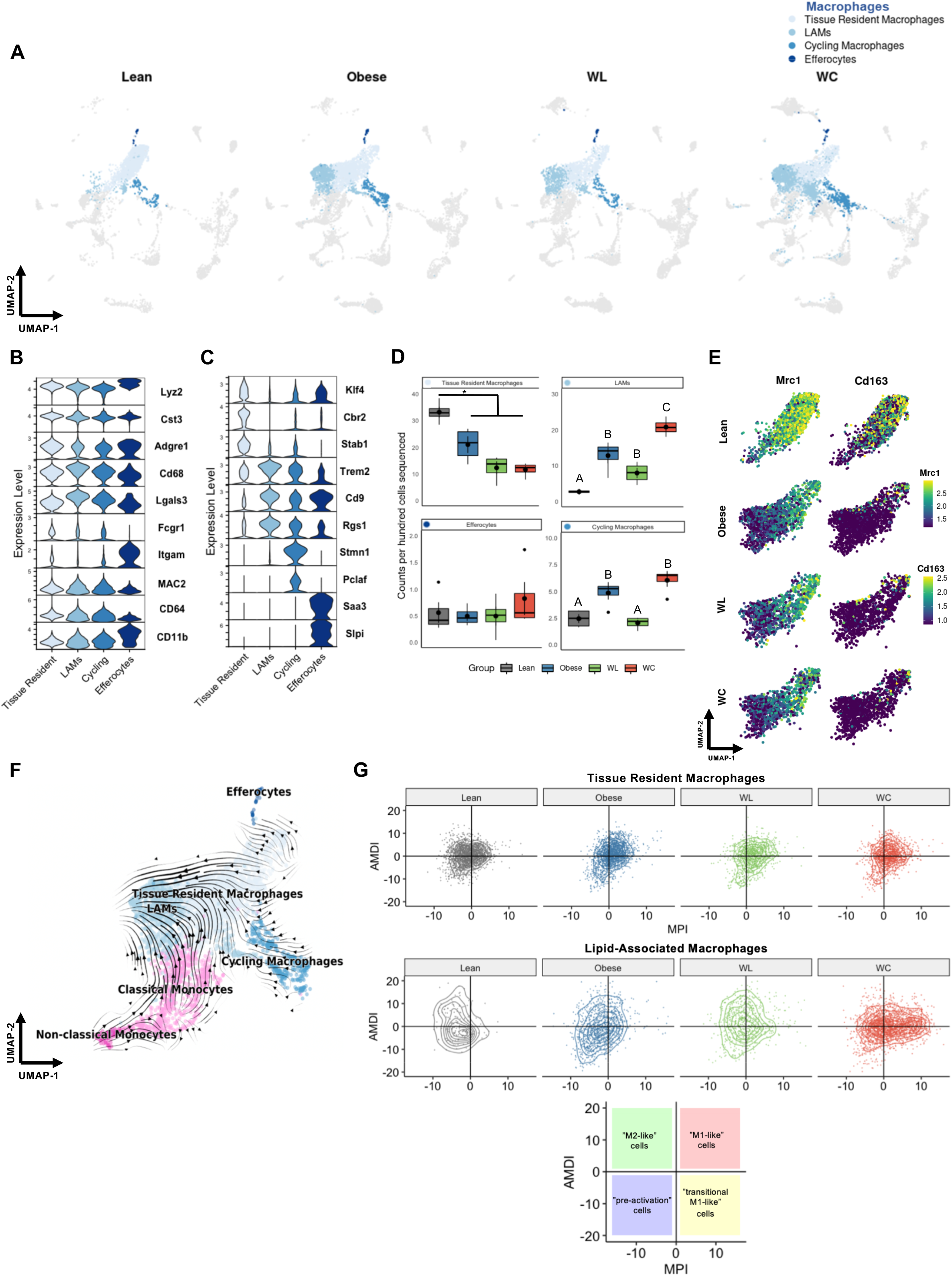
Diet-induced obesity causes persistent changes in adipose tissue macrophages. (A) Macrophage subclusters plotted by diet groups. (B) Expression levels for genes (*Lyz2, Cst3, Adgre1, Cd68, Lgals3, and Itgam*) and proteins (CITE-seq; MAC2, CD64, and CD11b) associated with macrophages. (C) Expression of genes that associate with macrophage subclusters. (D) Counts per hundred cells sequenced for macrophage subclusters (mean ± sem; n=4). E) UMAP visualization of Mrc1 and Cd163 expression in TRMs by diet group. (F) Embedding of RNA velocity displayed on the UMAP for macrophage and monocyte subsets. (G) TRMs and LAMs plotted on based on the macrophage polarization index (MPI) and the activation-induced macrophage differentiation index (AMDI) calculated using MacSpectrum. Pairwise two-tailed t-tests with Bonferroni correction for multiple comparisons were used to compare groups for cell counts (groups indicated by different letters are significantly different, \**adj p* < 0.05).

The changes in macrophage subclusters largely persist, even following 9 weeks of WL or WC. TRMs decreased with obesity and even more with WL and WC (**Fig 6D**). Importantly, while LAMs increase with obesity, they do not return to lean levels with WL and increase even more with WC; thus, levels are not directly correlated with AT mass. Cycling macrophages are increased with obesity, recovered with WL, but again worsen with WC and frequency of efferocytes was similar between diet groups.

### Alterations in macrophage phenotype remain unresolved with WL

A great challenge in classification of AT macrophages is that conventional markers are often not exclusive or only poorly describe cell function. We observed that TRMs retained high expression of the M2-associated Mrc1 gene (coding for the mannose receptor CD206). However, these same cells completely lost expression of the M2-associated Cd163 gene with the onset of obesity (**Fig 6E**). The persistence of this transcriptional change, which lasted for at least 9 weeks following WL and into subsequent WC, compelled us to further investigate if TRMs could be transitioning towards a LAM-like profile. Although the majority of LAMs are likely derived from tissue infiltrating monocytes, as previously suggested^18^, RNA velocity suggested that some TRMs are projected to become LAMs (**Fig 6F**). MacSpectrum, a tool which uses macrophage differentiation and polarization indexes previously generated using *in vitro* systems^17^, was used to further interrogate changes in tissue-resident and LAM phenotypes (**Fig 5G**). We observed that obesity shifted both subpopulations of macrophages towards a more pro-inflammatory phenotype that was not recovered following WL. Specifically, WC LAMs appeared to be highly inflammatory compared to cells from other conditions, indicating that these cells may be an important target for subsequent studies seeking to improve outcomes of WL and weight regain.

## Discussion

WL improves obesity-associated IR; however, as highlighted here and by others^8, 19, 28, 29^, WL does not normalize AT immune populations. Previous research shows that AT immune cells increase during early stages of WL^28^, presumably functioning as lipid handlers during lipolysis. In our model, animals maintain long term WL and still retain many of the changes associated with prior obesity. We postulate that lack of reversion of the immune phenotype to the lean state that occurs during WL, may ultimately be an underlying contributor to the detrimental impact of weight regain on cardiometabolic health. Supporting this, many inflammatory phenotypes are exacerbated in WC compared to the obese group. Indeed, given the frequency and risks associated with weight regain (**Fig 1** and ^7, 8, 9, 10, 11, 12, 13^), understanding the cell types and/or mechanisms in AT that are not fully recovered following WL is critical.

In the T cell compartment, the greatest change in abundance occurs in T_regs_, which decrease with obesity and do not rebound with WL. Interestingly, obese AT T_regs_ lose expression of *Il1rl,* which codes for the receptor ST2, and *Il1rl1* expression remains low with WL and WC. This is consistent with *Gata3* expressing ILC2s, which are reduced with obesity and produce IL33 – the primary ligand for ST2. Thus, AT T_reg_ maintenance may be impaired due to loss of IL-33 signaling, shown to promote glucose tolerance^30^. Interestingly, mast cells are increased with WC, but also have reduced *Il1lr* expression and increased expression of lipid-associated genes, such as *Trem2* and *Fabp5*. While the role of mast cells in AT is unclear due to lack of specific knockout models, this suggests that the demand to handle excess lipid in obese AT occurs concomitantly with reductions in availability and response to type 2 cytokines, such as IL-33, and that these changes are not resolved with WL.

CD8^+^ T cells are elevated in obesity and WC and are most enriched after WL. Moreover, obesity increases expression of exhaustion-associated genes in CD8^+^ T cells that is not normalized by WL. T cell exhaustion has been recently noted in human and mouse AT CD8^+^ T cells, which were shown to have impaired stimulation and increased markers of T cell exhaustion following obesity^31, 32, 33^. This transcriptional phenotype was further validated in our model by comparing T cells identified in our study to previously published exhausted T cell atlases generated in models of viral infection and cancer^24^. We also previously reported that AT T cell clonality is increased during obesity and that the T cell repertoire likely responds to positively charged, non-polar antigens^34^. It is plausible that APCs accumulate and present lipid-adducted proteins during lipid clearance following weight gain but also WL. Thus, chronic CD8+ T cell stimulation via antigen presentation may drive T cell exhaustion and warrants further investigation. In support of this, AT DCs shift to a mature, activated state with the onset of obesity and persist during WL and WC. This activated status is characterized by increased expression of *Ccr7, Mreg,* and *Fscn1*, which have all been previously reported as critical features of immunoregulatory DCs that are enriched in non-small-cell lung cancer and uptake of anti-tumor antigens^35^. Furthermore, expression of the immunoregulatory proteins *Cd274*, *Pdcd1lg2*, *and Cd200* on activated AT DCs suggests these cells may be important regulators of T cell function and potentially exhaustion in AT.

Macrophages are the most abundant immune cell type in our analysis. TRMs decrease with obesity and do not recover with subsequent WL. We also noticed a remarkable change in transcriptomic profile of this subcluster with obesity. Expression of *Mrc1*, coding for the M2-like macrophage marker CD206, is maintained during obesity, WL, and WC. However, expression of *Cd163*, another M2-like marker, is lost with the onset of obesity and the loss persists during WL and WC. These findings support divergence from the classical M1-M2 terminology, particularly in disease-associated microenvironments like obese AT, in favor of functional phenotyping or high-dimensional cell phenotyping^36^. Moreover, LAMs increase with obesity and only partly resolve with WL, supporting the notion that WL alone is insufficient to correct the AT immune landscape. Upon weight regain, LAMs and cycling macrophages tend to increase compared to obese animals. This shift in proportions of TRMs and LAMs suggests a critical role for lipid regulation in obesity, however, the role of macrophage lipid handling is not fully understood^37, 38^. LAM depletion via *Trem2* knockout impairs IR during HFD feeding, suggesting macrophages may help buffer lipid overload^18^. Additionally, loss of TREM2 via global knockout or anti-TREM2 neutralizing antibodies improves T cell responses, similar to anti-PD-1 immunotherapy, and ultimately reduced tumor size in a macrophage-dependent manner^39^. Thus, lipid handling may be delicately linked to antigen presentation and T cell exhaustion in both metabolic disease and cancer.

In summary, we identified three major immune activities that are induced by obesity, fail to resolve with WL, and are exacerbated in WC: 1) antigen presentation, 2) lipid handling, and 3) inflammation (**Fig 7**). Increases in activated DCs, LAMs, and exhausted T cells suggest that antigen presentation is a critical point of regulation, or dysregulation, in obese AT. Future studies will explore this with higher granularity by using single cell T cell receptor sequencing to uncover TCR clones associated with metabolic disease. Macrophage lipid handling also appears to be critical for AT regulation; however, whether this process reaches capacity or is dysregulated in obesity is not known. Moreover, dysregulation of macrophage lipid handling and the lipid scavenger receptor, TREM2, have been implicated as key features in numerous diseases, such as Alzheimer’s, cancer, and infection – expanding the importance of our findings to additional immunological diseases. Finally, the loss of type 2 immune subsets, IL-33 responsiveness, and changes in the TRM transcriptional phenotype along with a concomitant increase in M1-like cells, support an increase in inflammation, which has been previously linked to IR. Taken together, these findings support continued investigation into regulation of AT immune cells during obesity, WL, and WC. Studies utilizing other models of WL, such as exercise, bariatric surgery, and pharmacological WL, could greatly improve our understanding of immunometabolism and will help to identify advantageous immunotherapeutic targets.

**Figure 7.**
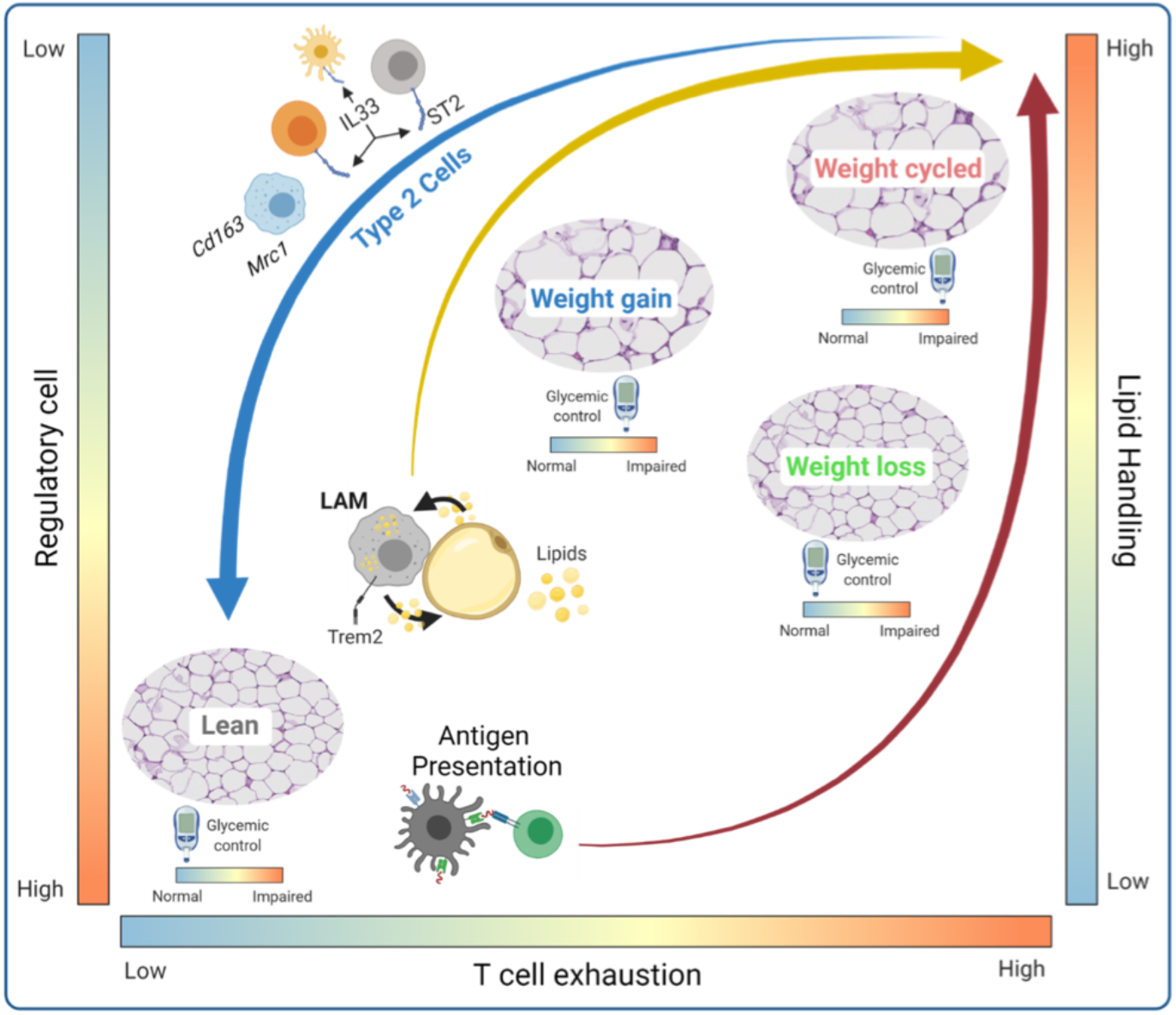
Adipose tissue immune remodeling during WL and WC. Body weight gain results in a transition from abundant type 2 immune cells present in lean adipose tissue towards an accumulation of immune cells/phenotypes associated with lipid handling (lipid associated macrophages, LAMs) and antigen presentation. WL does not restore the antigen presentation phenotype and is associated with T cell exhaustion. T cell exhaustion and LAMs are further amplified by weight regain. There is a disconnect between systemic glucose homeostasis and immune remodeling in adipose tissue, such that WL corrects obesity-induced glucose intolerance, but does not resolve the altered immune cell composition caused by diet-induced obesity.

The development of scSeq has been especially important in the field of Immunometabolism, as the dichotomy of immune cells as strictly type 1 or type 2 is inadequate and measuring changes in general cell populations mask changes that appear within cell type subclusters. We found 11 primary clusters and 33 unique subclusters, validated with CITE-seq antibodies for identifying cell surface markers. However, we did not capture all known AT immune populations. For example, we did not identify eosinophils, ILC1/3, B1-a and B2 cells, or iNKT cells in our dataset, which may be due to isolation methodology, inherent cell properties, and sequencing parameters. Eosinophils contain many RNAses for pathogen defense that are released during cell lysis. Moreover, it’s plausible that NKT and iNKT cells, for example, have differential cell surface expression that is not easily captured transcriptionally. Genes associated with transcription factors and cytokines were lowly expressed or dropped out in our data set, likely limiting our ability to identify cell types specifically distinguished by these markers. Use of stimulation conditions, increased sequencing depth, or CITE-seq targeted to intracellular epitopes^40^, could further improve identification of addition cell states in future experiments.

Importantly, we believe the immunophenotyping conducted using CITE-seq in our models of obesity, WL, and WC, has broad applicability, which can be hypothesis-generating for models of cancer, infection, or metabolic disease in other tissues. To facilitate discovery and to broaden accessibility of this data, we have created an open-access online interactive portal called MAIseq (Murine Adipose Immune sequencing) with our preprocessed data for the research community using modified source code from ShinyCell^41^ at https://hastylab.shinyapps.io/MAIseq/. Users can utilize a variety of built-in visualization approaches that span beyond what we could report in this manuscript. For instance, we also identified γ/δ T cells, NK and NKT cells, multiple B cell subsets and plasma cells (**Fig S8**), and even some CD45^+^ stromal cells that we did not investigate here. Users can explore our cell cluster annotations, identify potential genes of interest, plot gene expression by diet groups, clusters, or subclusters, and plot expression of surface markers from CITE-seq. We provide clear instructions for use and options to download figures and tables in a variety of formats. Collectively, our data provides critical groundwork for understanding the causes of WC-accelerated metabolic disease.

## Methods

### Mice Housing and Mouse Model

Male C57BL/6J mice were purchased from Jackson Labs at 7 wks of age. At 8 wks of age, mice were placed on 9 wk cycles of 60% HFD (Research Diets #D12492) or 10% LFD (Research Diets #D12450B) for a total of 27 weeks previously reported^14^ and shown in **Fig 1A**. Body weight and food intake were recorded weekly. The room was maintained at a controlled temperature (23°C) and 12 h light-dark cycles. All studies were approved by the Institutional Animal Care and Usage Committee of Vanderbilt University.

### Body Compensation and Glucose Tolerance

At 26 weeks, body composition was measured in the Vanderbilt University Mouse Metabolic Phenotyping Center via nuclear magnetic resonance (Bruker Minispec). Lean body mass, body fat, and free body fluid were recorded. The following day, ipGTT was performed. Briefly, baseline fasting blood samples were obtained by cutting off the tip of the tail under isoflurane anesthesia. Glucose was injected intraperitoneal at 1.5 g/kg of lean mass and subsequent blood samples were collected at 15, 30, 45, 60, 90, 120, and 150 min post injection. Blood glucose was measured using a Contour Next EZ Blood Glucose Monitoring System and test strips (Bayer). Differences in glucose tolerance over time were assessed using two-way ANOVAs and differences in body composition and glucose tolerance test AUC were assessed using one-way ANOVAs with Tukey post hoc analysis in Prism. Differences in tissue mass were assessed using a Wilcox Ranked-Sum test corrected for multiple comparisons. Statistical significance was set to adjusted *p* < 0.05.

### Immunofluorescence Microscopy

Tissues were fixed in 1% paraformaldehyde for one hour and stored in 70% EtOH overnight before paraffin embedding by the Vanderbilt Tissue Pathology Shared Resource core. Tissues were sectioned at 5 μm and allowed to dry overnight. Slides were deparaffinized in xylene and dehydrated through an EtOH gradient, then immunolabeled with Rabbit anti-Perilipin-1 (Abcam #9349T; Clone D1D8) at 1:200 overnight at 4 °C. After washing, Goat anti-Rabbit IgG conjugated to AF647 (Abcam #ab150079) was applied for two h at room temperature prior to washing and coverslipping with Prolong Gold (Invitrogen #P36931) containing DAPI. Slides were imaged using a 20X objective on a Leica DMI8 widefield microscope and captured with a Leica DFC9000GT camera. Image tiles were taken across the entirety of each section and stitched using the LAS X software suite. Merged images were processed in ImageJ and adipocytes were counted using an in-house macro. Briefly, the AF647 channel was processed by enhancing contrast and applying a gaussian blur with a sigma = 5. Background subtraction was performed with a rolling basis of 30. Auto thresholding using the “Triangle dark” setting was used to preserve adipocyte borders. The images were then skeletonized and then Image J’s analyze particles function was used with a size threshold range of 300-315000, circularity of 0.4-1.

### Stromal Vascular Fraction Isolation and Cell Sorting

Mice were euthanized by isoflurane overdose and cervical dislocation and perfused with 20 ml PBS through the left ventricle. eAT pads were collected, minced, and digested in 6 ml of 2-mg/mL type II collagenase (Worthington # LS004177) for 30 min at 37°C. Digested eAT was then vortexed, filtered through 100 μm filters, and lysed with ACK buffer, and filtered through 35 μm filters as previously described^42^.

The AT stromal vascular fraction was prepped with anti-mouse Fc Block (BD Biosciences). Cells from each mouse were labeled with unique hashtag antibodies (Biolegend TotalSeq-C) and anti-CD45 microbeads (Miltenyi). Biological replicates were pooled and sorted on a Miltenyi AutoMACs using the “possel_s” option. CD45+ cells were then labeled for surface markers using TotalSeq-C antibodies (Biolegend). All sample and antibody information can be found in **Supplemental Table 1**. Cells were stained with DAPI for FACS sorting and viable cells were collected for downstream processing and sequencing.

### Single Cell RNA-sequencing

All samples were submitted and processed for sequencing on the same day. Sample preparation was conducted using the 5’ assay for the 10X Chromium platform (10X Genomics) targeting 20,000 cells per diet group (~5,000 cells per biological replicate). 50,000 reads per cell were targeted for PE-150 sequencing on an Illumina NovaSeq6000. Sample processing and sequencing were completed in the VANderbilt Technologies for Advanced GEnomics core (VANTAGE).

### Data Processing

FastQ files obtained from sequencing were processed using CellRanger V3 (10X Genomics) with feature barcoding. Outputs from CellRanger were further processed using Velocyto^43^ for downstream RNA velocity analysis. The R package SoupX^44^ was used to remove ambient contaminating RNA. The Seurat V4 R package^45^ was used for quality control, data set integration, clustering, cell type annotation, differential expression, and visualization. The Python package scVelo^46^ was used for modeling and visualizing RNA velocity through R using Reticulate. For quality control of sequenced cells, only cells that had at least 200 gene features, at least 500 total measured RNA sequences, and less than 5% mitochondrial RNA content were retained. Furthermore, only cells designated as singlets after hashtag demultiplexing were used for visualization and analysis. RPCA-based integrated was used to integrate the four sequencing datasets. For more detailed processing information, we have provided the minimal necessary data and three vignettes containing code for preprocessing, data integration, and cell type annotation at https://github.com/HastyLab/Multiomics-WeightCycling-Vignettes.

### Annotating clusters

Cluster annotation was performed following dimensional reduction using the integrated data assay. First, broad clusters were identified using low resolution (0.4) and annotated by common cell type markers. Each identified broad cluster, referred to as “Cell Types”, were further subclustered and subclusters were annotated by comparing differentially expressed markers to published literature. Cluster annotations were further supported using SingleR with the Immgen and MouseRNAseq databases^47^.

### Downstream sequencing analysis

All downstream analyses comparing cells across diet groups, individual biological replicates, or clusters were performed using the normalized RNA assay. Differential expression was performed using the Wilcoxon Ranked-Sums test in Seurat V4 for single cells and the likelihood ratio test in DESeq2 for pseudobulk data. Pathway analysis was conducted on pseudobulk data using Metascape^48^. Differential abundance testing was conducted using the MiloR package^49^ in R. Metaclusters identified by MiloR were further tested with permutation testing using the scProportionTest R package (https://github.com/rpolicastro/scProportionTest). For exploring macrophage phenotype, the R application MacSpectrum^17^ was used with provided index information.

## Data Availability

Raw sequencing files, processed data matrices, UMAP and PCA embeddings, cell metadata, and a fully integrated Seurat v4 R data object are available via the NCBI GEO with the primary accession code GSE182233.

## Code Availability

Example code to reproduce our processed and integrated Seurat v4 object is available at: https://github.com/HastyLab/Multiomics-WeightCycling-Vignettes. This GitHub repository contains information for 1) Preprocessing and Quality Control, 2) Data Integration, and 3) Cell Type Annotation.

All code necessary to reproduce the figures within this manuscript are available at via GitHub at: https://github.com/HastyLab/Multiomics-WeightCycling-Figures.

Code for generating our ShinyCell-based web application is available upon request, but we highly recommend installation of ShinyCell directly through the author’s GitHub page.

## Acknowledgements

This project was funded by a Veterans Affairs Merit Award 5I01BX002195 and an AHA Innovation Award (19IPLOI4760376) to AHH. MAC is funded by an NIH F31 Predoctoral Fellowship (1F31DK123881), HLC is funded by an AHA Postdoctoral Fellowship (20POST35120547), and NCW is funded by an AHA Postdoctoral Fellowship (21POST834990). MAC, HLC, and NCW were all previously supported by the Molecular Endocrinology Training Program (T32 DK07563). The Translational Pathology Shared Resource used for tissue preparation is supported by NCI/NIH Cancer Center Support Grant 5P30 CA68485-19. FACS sorting was performed in the Vanderbilt Flow Cytometry Shared Resource which is supported by the Vanderbilt Ingram Cancer Center (P30CA068485) and the Vanderbilt Digestive Disease Research Center (DK058404). Single-cell preparation and sequencing were performed in the Vanderbilt Technologies for Advanced Genomics (VANTAGE) core laboratory. Figures 1A, 2A, and 7 were produced with BioRender.com.

## Author Contributions

MAC and HLC completed the mouse work and cell isolation and processing for these experiments. MAC performed data analysis and generated the interactive web portal. HLC and MAC drafted the manuscript. MAC and NCW produced the experimental design and summary figures using BioRender.com. AHH conceptualized the study, provided funding, and is the guarantor for this work. All authors contributed to experimental design, data interpretation, website testing, and manuscript revisions.

---

**Supplemental Figure 1.**
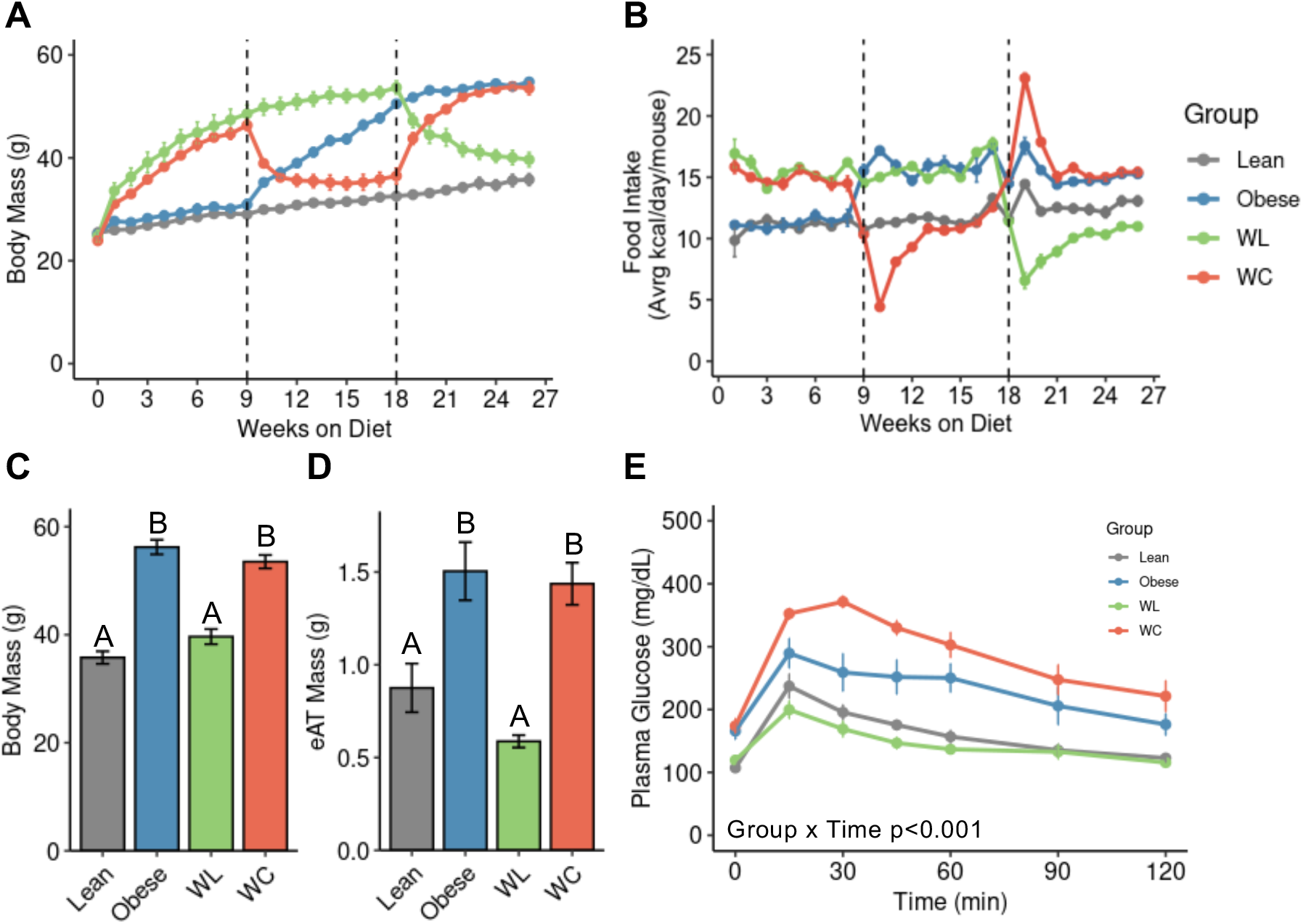
Body mass and glucose tolerance for mice used for CITEseq analysis. (A) Body mass over time measured weekly. (B) Food intake over time measured weekly. (C) Body mass and (D) eAT mass measured at sacrifice. (E) Intraperitoneal glucose tolerance test dosed at 1.5 g dextrose/kg lean mass one week prior to sacrifice. Pairwise two-tailed T-tests with Bonferroni correction for multiple comparisons were used to compare groups for body mass and eAT mass and two-way ANOVA was used to compare groups for ipGTT (groups indicated by different letters are significantly different, *adj p* < 0.05). Data are plotted as mean ± SEM (n=4).

**Supplemental Figure 2.**
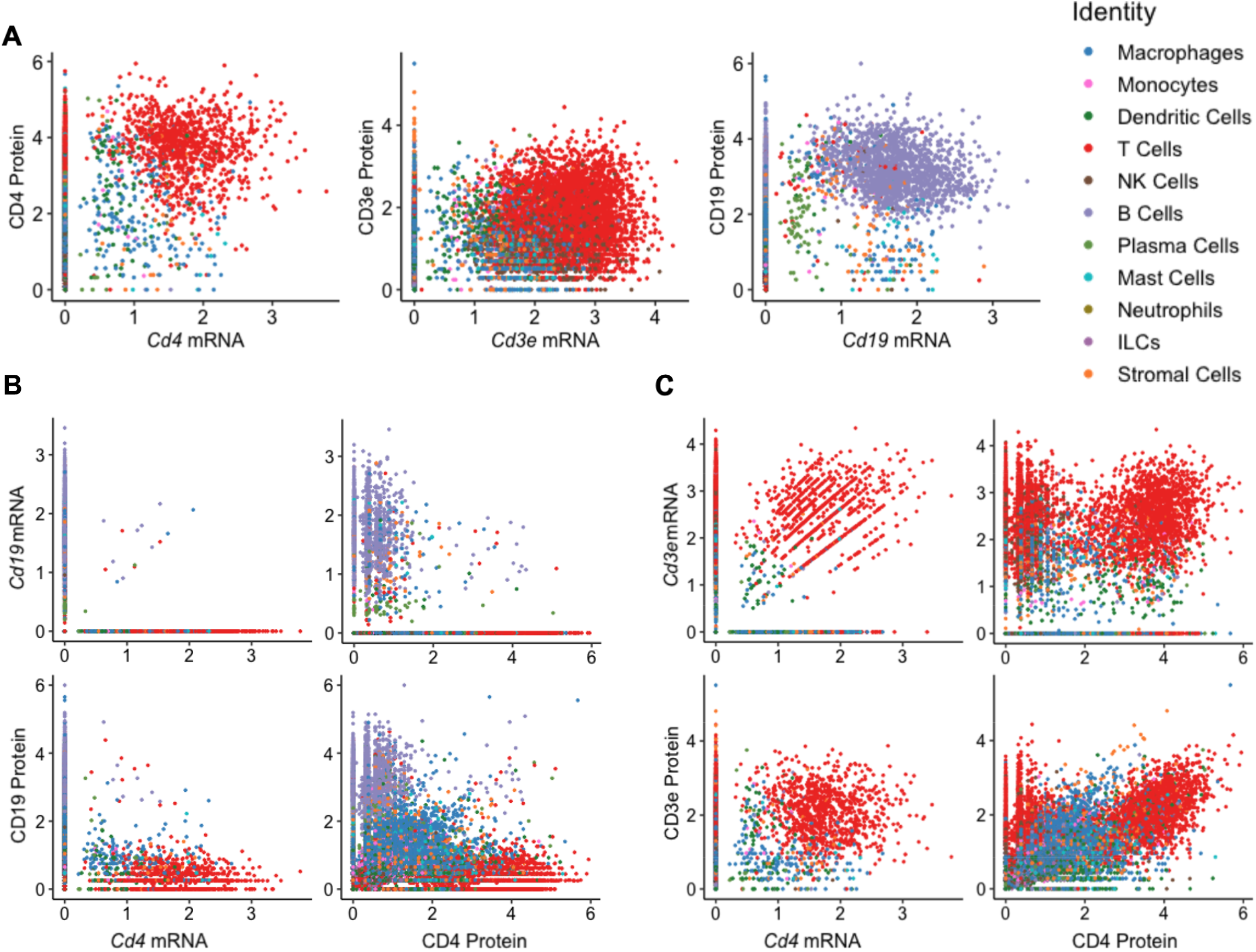
Validation of CITE-seq antibodies. (A) Comparison of genes for *Cd3e, and Cd19* with their corresponding surface proteins measured using CITE-seq. Individual cells are colored by annotated cell type. (B) Comparison of cell type exclusive genes *Cd4* and *Cd19* with their corresponding surface proteins. (C) Comparison of co-expressed genes *Cd4* and *Cd3e* with their corresponding surface proteins.

**Supplemental Figure 3.**
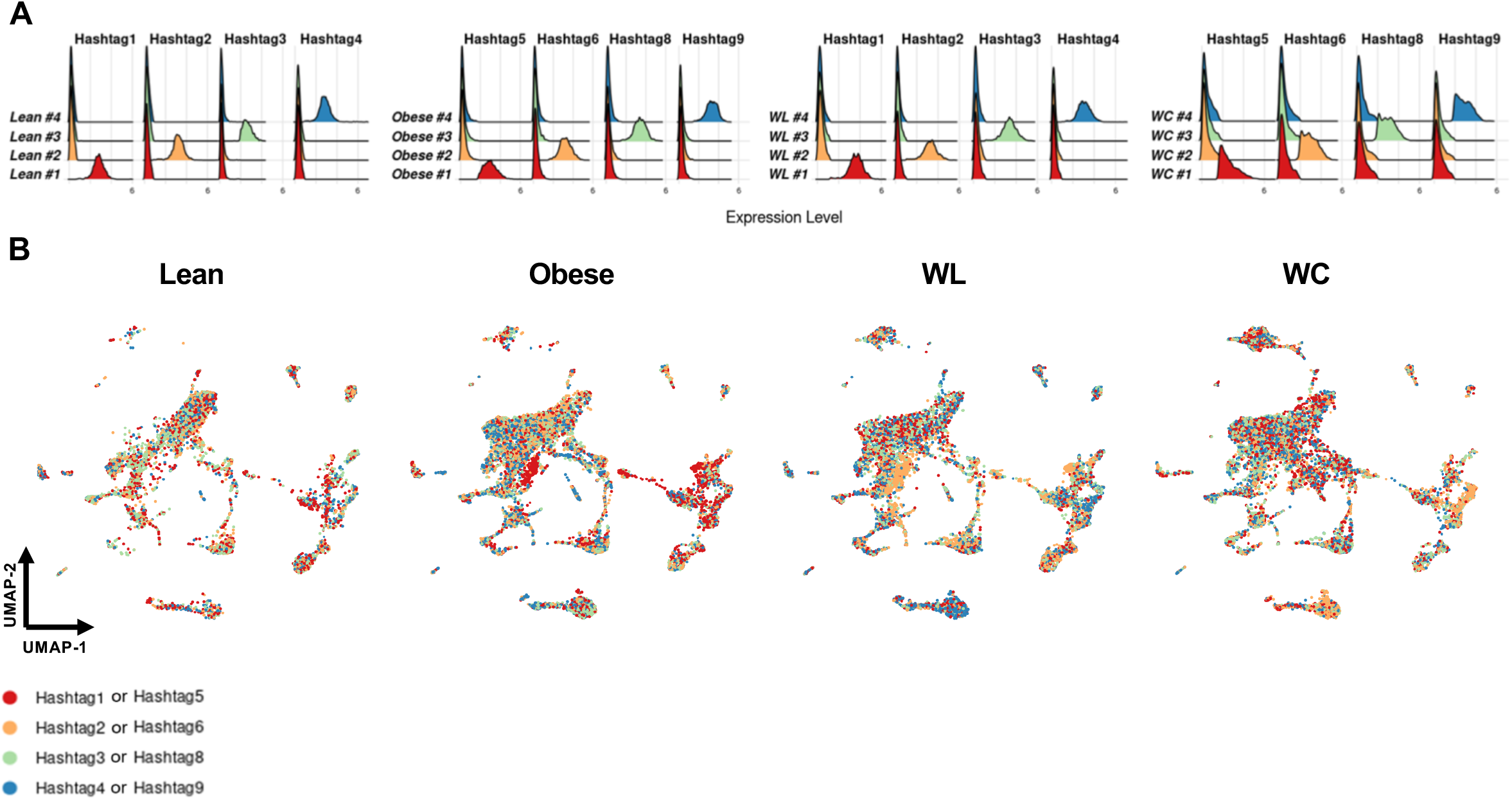
Biological replicates identified by Hashtag demultiplexing are well represented within diet groups. (A) Hashtags demultiplexed for each individual mouse per diet group (n=4 per group; total of 16 mice sequenced). (B) Uniform manifold projection colored by Hashtag for each diet group.

**Supplemental Figure 4.**
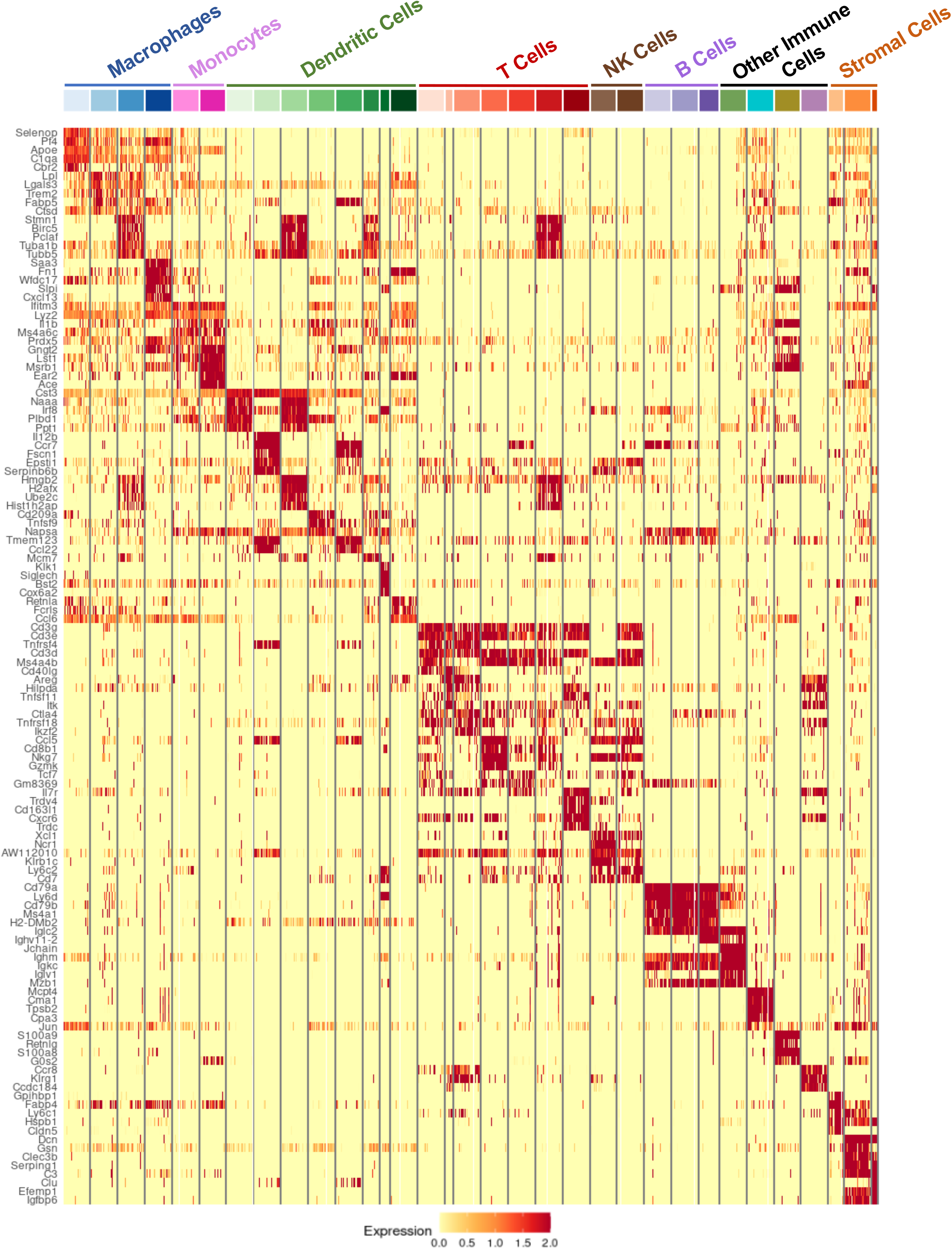
Heatmap of top five differentially expressed genes for each high-resolution cluster identity. Each row is a unique gene, colored red when expressed highly. Each thin vertical line indicates one subsampled cell (up to 100 cells per subcluster).

**Supplemental Figure 5.**
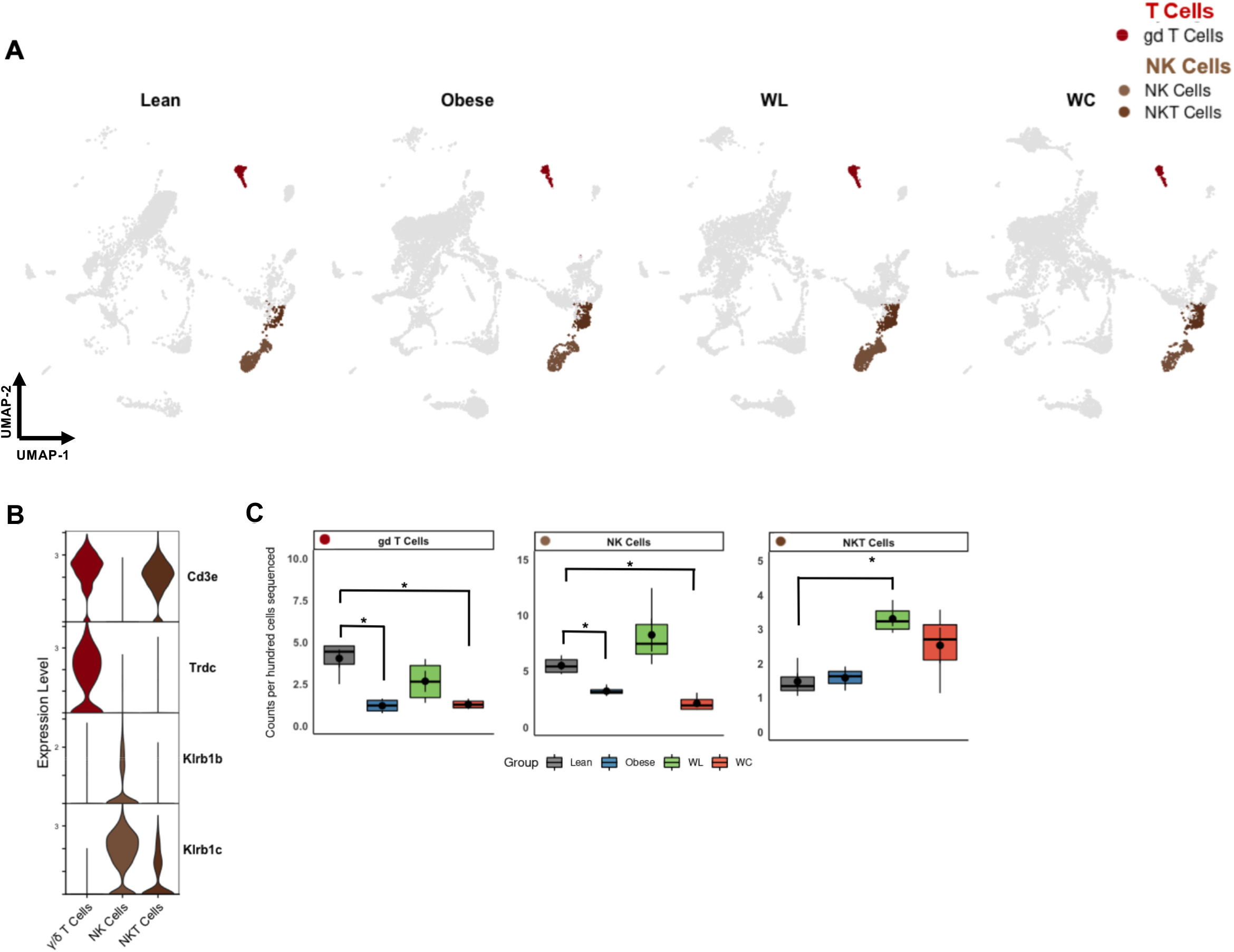
Frequency and markers of adipose tissue γ/δ, NK, and NKT cells. (A) UMAP of γ/δ T cells and NK cell subclusters by diet group. (B) Expression of markers enriched in γ/δ T cells and NK cell subclusters. (C) Counts per hundred cells sequenced for γ/δ T cells and NK cell subclusters (mean ± SEM; n=4). Pairwise two-tailed t-tests with Bonferroni correction for multiple comparisons were used to compare groups for cell counts (*adj *p* < 0.05).

**Supplemental Figure 6.**
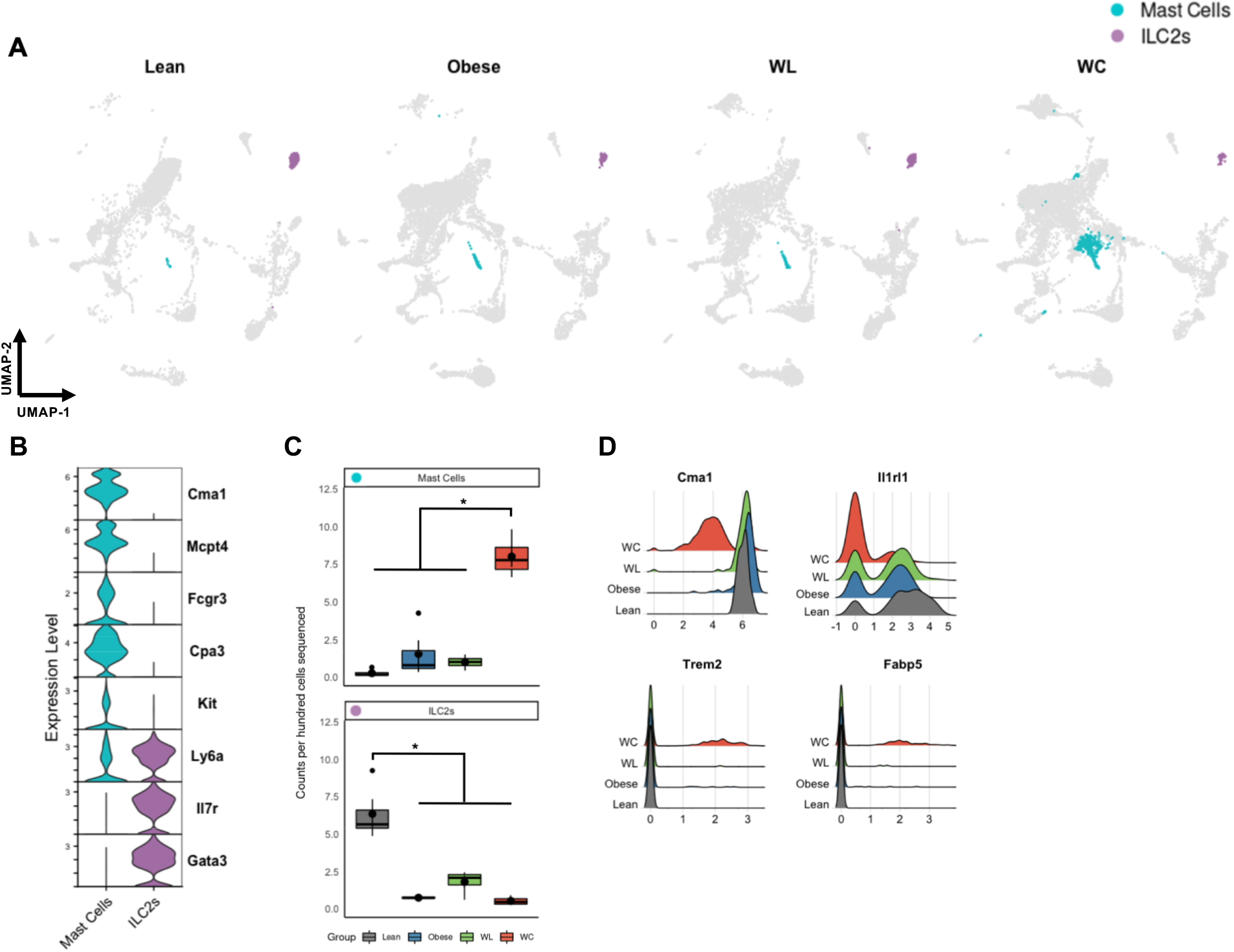
Adipose tissue mast cells have reduced expression of *Ilr1l1* and ILC2s are reduced in frequency after obesity. (A) UMAP of mast cells and ILC2s by diet group. (B) Expression of markers enriched in mast cells and ILC2 subclusters. (C) Counts per hundred cells sequenced for mast cells and ILC2s (mean ± sem; n=4). (D) Expression of *Cma1* coding for the secreted protease chymase, *Il1rl1* coding for the IL33-R, and lipid-associated proteins *Trem2*, and *Fabp5* in mast cells by diet group. Pairwise two-tailed t-tests with Bonferroni correction for multiple comparisons were used to compare groups for cell counts (*adj *p* < 0.05).

**Supplemental Figure 7.**
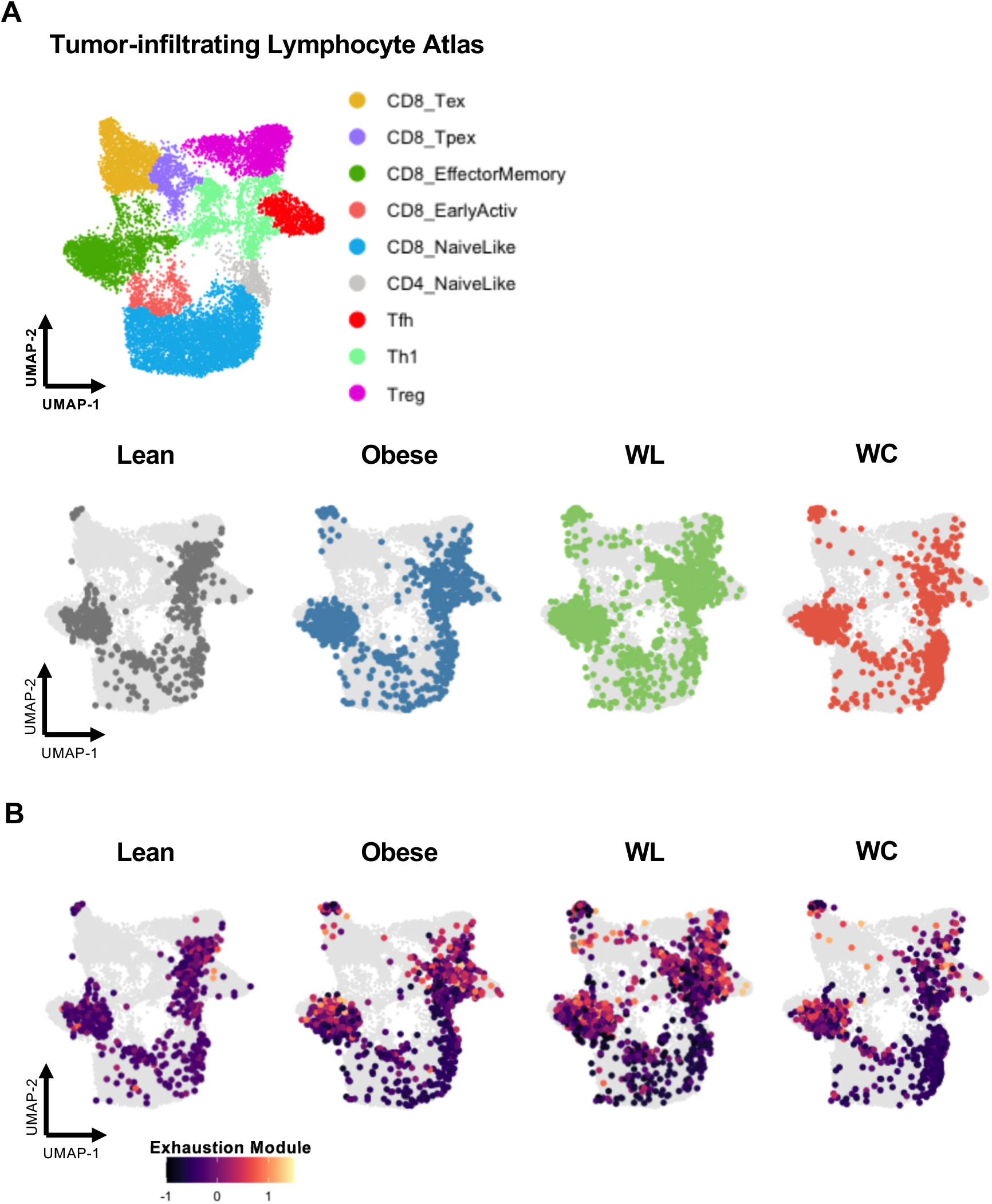
Alignment of CD8+ T cells to a tumor-infiltrating lymphocyte reference atlas. (A) T cells separated by diet group plotted onto a tumor-infiltrating lymphocyte reference atlas using ProjecTILs. (B) T cells, projected onto the TIL reference atlas, colored by an exhaustion module containing the features *Pdcd1, Tox, Tigit, Lag3, and Entpd1*. Note that all though the reference atlas suggests the majority of cells are effector memory, they express high levels of genes within the T cell exhaustion module.

**Supplemental Figure 8.**
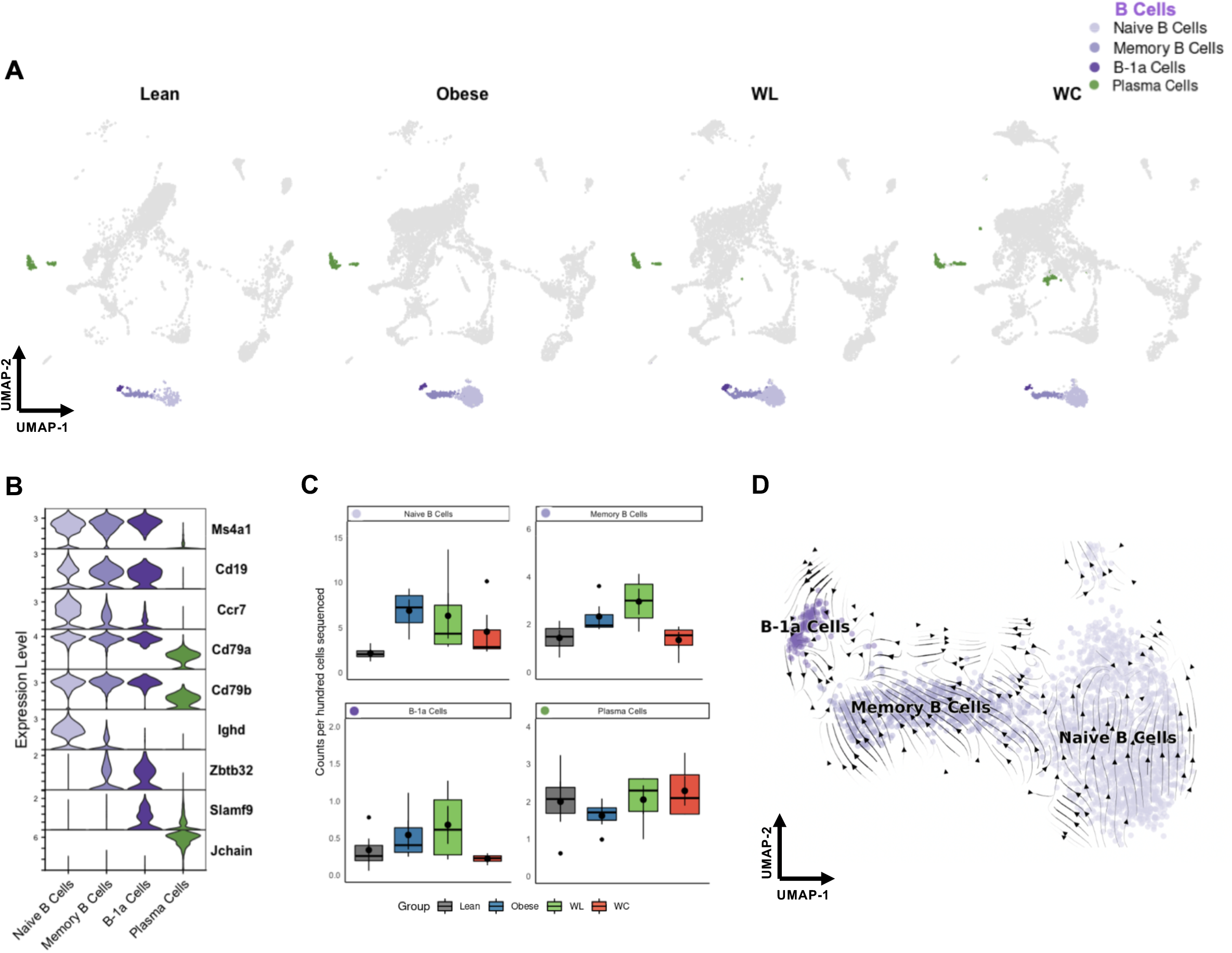
Frequency and markers of adipose tissue B and plasma cells. (A) UMAP of B cell subclusters and plasma cells by diet group. (B) Expression of markers enriched in B cell subclusters and plasma cells. (C) Counts per hundred cells sequenced for B cell subclusters and plasma cells (mean ± SEM; n=4). (D) Embedding of RNA velocity displayed on the UMAP for B cell subsets. Pairwise two-tailed t-tests with Bonferroni correction for multiple comparisons were used to compare groups for cell counts (no significant differences observed for *adj p* < 0.05).

## References

1. Hruby, A. & Hu, F.B. The Epidemiology of Obesity: A Big Picture. Pharmacoeconomics 33, 673–689 (2015).

2. Organization, W.H. Obesity and overweight. 2021 [cited]Available from: https://www.who.int/news-room/fact-sheets/detail/obesity-and-overweight.

3. Ferrante, A.W., Jr. Macrophages, fat, and the emergence of immunometabolism. J Clin Invest 123, 4992–4993 (2013).

4. Russo, L. & Lumeng, C.N. Properties and functions of adipose tissue macrophages in obesity. Immunology 155, 407–417 (2018).

5. Zatterale, F. et al. Chronic Adipose Tissue Inflammation Linking Obesity to Insulin Resistance and Type 2 Diabetes. Front Physiol 10, 1607 (2019).

6. Khan, S., Chan, Y.T., Revelo, X.S. & Winer, D.A. The Immune Landscape of Visceral Adipose Tissue During Obesity and Aging. Front Endocrinol (Lausanne) 11, 267 (2020).

7. Fildes, A. et al. Probability of an Obese Person Attaining Normal Body Weight: Cohort Study Using Electronic Health Records. Am J Public Health 105, e54–59 (2015).

8. Wing, R.R.P. S. Long-term weight loss maintenance. Am J Clin Nutr 82, 222S–225S (2005).

9. Crawford, D.J., R.W.; French, S.A. Can anyone successfully control their weight? Findings of a three year community-based study of men and women. Int J Obes Relat Metab Disord 24, 1107–1110 (2000).

10. Stunkard, A. The results of a treatment for obesity: a review of the literature and report of a series. AMA Arch. Intern. Med. 108 (1959).

11. Delahanty, L.M. et al. Effects of weight loss, weight cycling, and weight loss maintenance on diabetes incidence and change in cardiometabolic traits in the Diabetes Prevention Program. Diabetes Care 37, 2738–2745 (2014).

12. Bangalore, S. et al. Body-Weight Fluctuations and Outcomes in Coronary Disease. N Engl J Med 376, 1332–1340 (2017).

13. Rzehak, P. et al. Weight change, weight cycling and mortality in the ERFORT Male Cohort Study. Eur J Epidemiol 22, 665–673 (2007).

14. Anderson, E.K., Gutierrez, D.A., Kennedy, A. & Hasty, A.H. Weight cycling increases T-cell accumulation in adipose tissue and impairs systemic glucose tolerance. Diabetes 62, 3180–3188 (2013).

15. Zou, J. et al. CD4+ T cells memorize obesity and promote weight regain. Cell Mol Immunol 15, 630–639 (2018).

16. Kratz, M. et al. Metabolic dysfunction drives a mechanistically distinct proinflammatory phenotype in adipose tissue macrophages. Cell Metab 20, 614–625 (2014).

17. Li, C. et al. Single cell transcriptomics based-MacSpectrum reveals novel macrophage activation signatures in diseases. JCI Insight 5 (2019).

18. Jaitin, D.A. et al. Lipid-Associated Macrophages Control Metabolic Homeostasis in a Trem2-Dependent Manner. Cell 178, 686–698 e614 (2019).

19. Weinstock, A. et al. Single-Cell RNA Sequencing of Visceral Adipose Tissue Leukocytes Reveals that Caloric Restriction Following Obesity Promotes the Accumulation of a Distinct Macrophage Population with Features of Phagocytic Cells. Immunometabolism 1 (2019).

20. Stoeckius, M. et al. Simultaneous epitope and transcriptome measurement in single cells. Nat Methods 14, 865–868 (2017).

21. Stoeckius, M. et al. Cell Hashing with barcoded antibodies enables multiplexing and doublet detection for single cell genomics. Genome Biol 19, 224 (2018).

22. Weinstock, A., Moura Silva, H., Moore, K.J., Schmidt, A.M. & Fisher, E.A. Leukocyte Heterogeneity in Adipose Tissue, Including in Obesity. Circ Res 126, 1590–1612 (2020).

23. Kane, H. & Lynch, L. Innate Immune Control of Adipose Tissue Homeostasis. Trends Immunol 40, 857–872 (2019).

24. Andreatta, M. et al. Projecting single-cell transcriptomics data onto a reference T cell atlas to interpret immune responses. Preprint at https://www.biorxiv.org/content/10.1101/2020.06.23.166546v1 (2020).

25. Orecchioni, M., Ghosheh, Y., Pramod, A.B. & Ley, K. Macrophage Polarization: Different Gene Signatures in M1(LPS+) vs. Classically and M2(LPS-) vs. Alternatively Activated Macrophages. Front Immunol 10, 1084 (2019).

26. Stonier, S.W., Ma, L.J., Castillo, E.F. & Schluns, K.S. Dendritic cells drive memory CD8 T-cell homeostasis via IL-15 transpresentation. Blood 112, 4546–4554 (2008).

27. Roberts, A.W. et al. Tissue-Resident Macrophages Are Locally Programmed for Silent Clearance of Apoptotic Cells. Immunity 47, 913–927 e916 (2017).

28. Zamarron, B.F. et al. Macrophage Proliferation Sustains Adipose Tissue Inflammation in Formerly Obese Mice. Diabetes 66, 392–406 (2017).

29. Vatarescu, M. et al. Adipose tissue supports normalization of macrophage and liver lipid handling in obesity reversal. J Endocrinol 233, 293–305 (2017).

30. de Oliveira, M.F.A., Talvani, A. & Rocha-Vieira, E. IL-33 in obesity: where do we go from here? Inflamm Res 68, 185–194 (2019).

31. Porsche, C.E., Delproposto, J.B., Geletka, L., O’Rourke, R. & Lumeng, C.N. Obesity results in adipose tissue T cell exhaustion. JCI Insight 6 (2021).

32. Shirakawa, K. et al. Obesity accelerates T cell senescence in murine visceral adipose tissue. J Clin Invest 126, 4626–4639 (2016).

33. Shirakawa, K. et al. Negative legacy of obesity. PLoS One 12, e0186303 (2017).

34. McDonnell, W.J. et al. High CD8 T-Cell Receptor Clonality and Altered CDR3 Properties Are Associated With Elevated Isolevuglandins in Adipose Tissue During Diet-Induced Obesity. Diabetes 67, 2361–2376 (2018).

35. Maier, B. et al. A conserved dendritic-cell regulatory program limits antitumour immunity. Nature 580, 257–262 (2020).

36. Felix, I. et al. Single-Cell Proteomics Reveals the Defined Heterogeneity of Resident Macrophages in White Adipose Tissue. Front Immunol 12, 719979 (2021).

37. Xu, X. et al. Obesity activates a program of lysosomal-dependent lipid metabolism in adipose tissue macrophages independently of classic activation. Cell Metab 18, 816–830 (2013).

38. Coats, B.R. et al. Metabolically Activated Adipose Tissue Macrophages Perform Detrimental and Beneficial Functions during Diet-Induced Obesity. Cell Rep 20, 3149–3161 (2017).

39. Molgora, M. et al. TREM2 Modulation Remodels the Tumor Myeloid Landscape Enhancing Anti-PD-1 Immunotherapy. Cell 182, 886–900 e817 (2020).

40. Chung, H. et al. Simultaneous single cell measurements of intranuclear proteins and gene expression. Preprint at https://www.biorxiv.org/content/10.1101/2021.01.18.427139v1 (2021).

41. Ouyang, J.F., Kamaraj, U.S., Cao, E.Y. & Rackham, O.J.L. ShinyCell: Simple and sharable visualisation of single-cell gene expression data. Bioinformatics (2021).

42. Orr, J.S., Kennedy, A.J. & Hasty, A.H. Isolation of adipose tissue immune cells. J Vis Exp, e50707 (2013).

43. La Manno, G. et al. RNA velocity of single cells. Nature 560, 494–498 (2018).

44. Young, M.D. & Behjati, S. SoupX removes ambient RNA contamination from droplet-based single-cell RNA sequencing data. Gigascience 9 (2020).

45. Hao, Y. et al. Integrated analysis of multimodal single-cell data. Cell 184, 3573–3587 e3529 (2021).

46. Bergen, V., Lange, M., Peidli, S., Wolf, F.A. & Theis, F.J. Generalizing RNA velocity to transient cell states through dynamical modeling. Nat Biotechnol 38, 1408–1414 (2020).

47. Aran, D. et al. Reference-based analysis of lung single-cell sequencing reveals a transitional profibrotic macrophage. Nat Immunol 20, 163–172 (2019).

48. Zhou, Y. et al. Metascape provides a biologist-oriented resource for the analysis of systems-level datasets. Nat Commun 10, 1523 (2019).

49. Dann, E., Henderson, N.C., Teichmann, S.A., Morgan, M.D. & Marioni, J.C. Milo: differential abundance testing on single-cell data 2 using k-NN graphs. Preprint at https://www.biorxiv.org/content/10.1101/2020.11.23.393769v1 (2020).

