## Supplemental Table 1 for "Multiomics reveals persistence of obesity-associated immune cell phenotypes in adipose tissue during weight loss and subsequent weight regain"

| Mouse Information | Hashing Antibody | Pooled Sample ID |
|---|---|---|
| Mouse 1, Lean | 0301 anti-mouse Hashtag 1 | 1 |
| Mouse 2, Lean | 0302 anti-mouse Hashtag 2 | 1 |
| Mouse 3, Lean | 0303 anti-mouse Hashtag 3 | 1 |
| Mouse 4, Lean | 0304 anti-mouse Hashtag 4 | 1 |
| Mouse 5, Obese | 0305 anti-mouse Hashtag 5 | 2 |
| Mouse 6, Obese | 0306 anti-mouse Hashtag 6 | 2 |
| Mouse 7, Obese | 0308 anti-mouse Hashtag 8 | 2 |
| Mouse 8, Obese | 0309 anti-mouse Hashtag 9 | 2 |
| Mouse 9, WL | 0301 anti-mouse Hashtag 1 | 3 |
| Mouse 10, WL | 0302 anti-mouse Hashtag 2 | 3 |
| Mouse 11, WL | 0303 anti-mouse Hashtag 3 | 3 |
| Mouse 12, WL | 0304 anti-mouse Hashtag 4 | 3 |
| Mouse 13, WC | 0305 anti-mouse Hashtag 5 | 4 |
| Mouse 14, WC | 0306 anti-mouse Hashtag 6 | 4 |
| Mouse 15, WC | 0308 anti-mouse Hashtag 8 | 4 |
| Mouse 16, WC | 0309 anti-mouse Hashtag 9 | 4 |

| Cell Sorting Reagents | Manufacturer | Catalog |
|---|---|---|
| Fc Block | BD Biosciences | 553142 |
| CD45 microbeads | Mitenyi | 130-052-301 |
| DAPI | Thermofischer | 62247 |

| TotalSeq-C Feature Barcoding Antibodies | Manufacturer | Catalog |
|---|---|---|
| anti-mouse MAC2/GAL3 | Biolegend | 125423 |
| anti-mouse CD279/PD-1 | Biolegend | 109127 |
| anti-mouse FCγR1 | Biolegend | 139327 |
| anti-mouse CD4 | Biolegend | 100571 |
| anti-mouse CCR7/CD197 | Biolegend | 120131 |
| anti-mouse CD80 | Biolegend | 109127 |
| anti-mouse CD11c | Biolegend | 117361 |
| anti-mouse CD44 | Biolegend | 103063 |
| anti-mouse NK1.1 | Biolegend | 108765 |
| anti-mouse TCRγ/δ | Biolegend | 118141 |
| anti-mouse CD39 | Biolegend | 143815 |
| anti-mouse CD19 | Biolegend | 115571 |
| anti-mouse CD11b | Biolegend | 101275 |
| anti-mouse CD3 | Biolegend | 100263 |
| anti-mouse TIGIT | Biolegend | 142119 |
| anti-mouse CD8α | Biolegend | 100785 |
